# Spatial organization of multisensory convergence in mouse isocortex

**DOI:** 10.1101/2024.12.09.627642

**Authors:** Kinjal Patel, Avery Hee-Woon Ryoo, Michael Buice, Stefan Mihalas, Bryan Tripp

## Abstract

The diverse functions of different cortical areas are thought to arise from their distinct groups of inputs. However, additional organizing principles may exist in the spatial structure of converging inputs. We investigated spatial convergence patterns of projections from primary sensory areas to other areas throughout the mouse isocortex. We used a large tract tracing dataset to estimate the dimension of the space into which topographical connections from multiple modalities converged within each other cortical area. We call this measure the topography dimension (TD). TD is higher for areas that receive inputs of similar strength from multiple sensory modalities, and lower when multiple inputs terminate in register with one another. Across the isocortex, TD varied by a factor of 4. TD was positively correlated with hierarchy score, an independent measure that is based on laminar connection patterns. Furthermore, TD (an anatomical measure) was significantly related to several measures of neural activity. In particular, higher TD was associated with higher neural activity dimension, lower population sparseness, and lower lifetime sparseness of spontaneous activity, independent of an area’s hierarchical position. Finally, we analyzed factors that limited TD and found that linear correlations among projections from different areas typically had little impact, while diversity of connection strengths, both between different projections onto the same area, and within projections across different parts of an area, limited TD substantially. This analysis revealed additional intricacy of cortical networks, beyond areas’ sets of connections and hierarchical organization. We propose a means of approximating this organization in deep-network models.

## 2 Introduction

Multisensory information is merged at multiple levels, including peripherally [8], subcortically [67, 52, 4], in primary sensory areas [17, 41, 2, 39, 29, 15, 46], and extensively in higher cortical areas [33, 45, 57, 65]. In humans, multisensory integration affects the categorization of objects [7] and events [34]. It can improve the detection of weak signals [35], make perceptual estimates more accurate [11, 49], resolve sensory ambiguities [20], and shorten reaction times [24]. Multisensory integration also provides a basis for responding to events when one modality is attenuated [13]. In artificial networks, multimodal convergence can provide a basis for self-supervised learning [47, 50].

Multisensory integration depends on the physical convergence of multisensory signals. The anatomy of multisensory convergence has been extensively studied in terms of the existence and strength of connections between brain structures. However, the integration of multisensory information by individual neurons also depends on the topography of these connections [25]. In the superior colliculus, visual, auditory, and tactile signals are organized topographically and aligned with each other [61]. The primate parietal cortex also contains converging topographic maps of multiple modalities [56]. For example, the parietal cortex of both human [22] and macaque [10] contains regions that respond to both tactile stimuli and visual looming toward the same receptive fields. These examples notwithstanding, the topography of multisensory convergence has generally received less attention than the existence of such connections. For example, a recent study [18] investigated inputs to mouse multisensory areas, however, tracer injections were always made into the same coordinates in each area, precluding investigation of connection topography.

Detailed data on multisensory topography is implicitly available in public tract tracing datasets, but organizing principles are needed to make sense of this large volume of data. A fundamental property of the confluence of multiple maps is the dimension of the space they are mapped into. To describe this property, we introduce the topography dimension (TD). TD is calculated by applying dimensionality reduction to a matrix of topographical coordinates representing multiple sensory modalities across different locations within a brain structure. For a given structure, the TD is lower if it receives projections from fewer sensory modalities, or if they converge in register (as in superior colliculus). It is highest when many sensory modalities project onto the structure with similar strength and diverse spatial patterns. We estimated TD for all neocortical areas of the mouse, and found that it varied across the neocortex by a factor of roughly four. We examined the relationship between TD of mouse neocortical areas and their hierarchy score [21], and found a significant positive correlation. We also investigated whether TD (a structural measure) is related to functional measures (including spontaneous neural activity dimension, population sparseness, and temporal sparseness), and found significant relationships in each case. TD improved prediction of each of these measures, even after accounting for HS, and it was more consistently related than HS in different analyses (e.g. standard vs. robust regression).

Anchoring functional mouse brain studies into the Allen Mouse Brain Common Coordinate Framework (CCF) [68] has become a standard in the field [e.g. 59, 32]. The atlas contains functional associations in its naming of sensory areas, such as VISp for primary visual area, or VISrl for rostrolateral visual area. However, these simple functional names hide a more complex multisensory integration. For example, VISrl contains a lot of somatosensory input. We believe the analysis preformed in this study, which is integrated in the CCF, will allow follow-up functional studies to better relate to the areas’ complex sensory inputs.

As far as we are aware, this work is the first systematic characterization of the structure of multisensory convergence throughout the neocortex and reveals the strongest quantitative predictions so far of fundamental functional measures from cortical structure.

## 3 Results

### 3.1 Topography dimension across the isocortex

We analysed the structure of connections from six primary sensory areas: The primary visual and auditory areas (VISp and AUDp) and four primary somatosensory areas, specifically the barrel field (SSp-bfd), upper limb (SSp-ul), nose (SSp-n), and mouth (SSp-m) areas.

We studied multisensory convergence using the mesoscale connectivity model of [30]. This is a voxel-to-voxel map of connection density that is based on 428 anterograde tract tracing experiments from the Allen Mouse Brain Connectivity Atlas [44]. Based on these experiments, [30] used kernel regression to estimate connection weights between 100 *µ*m voxels throughout the mouse brain.

We assigned 2D cortical-surface coordinates to each primary-area voxel. We then propagated these coordinates through the mesoscale model and found the weighted-average input coordinates to each other voxel in the isocortex. This resulted in 12D coordinates for each voxel (i.e., weighted averages of two cortical-surface dimensions from each of six primary sensory areas).

For each isocortical area, we assembled a matrix of coordinates with twelve rows and one column per voxel in that area. We multiplied these coordinates by the weighted sums of corresponding voxel-to-voxel connection weights to compress the coordinates of weak connections. We then calculated the dimension of the resulting matrix as the participation ratio [14] of the eigenvalues of its correlation matrix. We call this measure the topography dimension (TD) of the area. Across the mouse isocortex, TD based on inputs from VISp, AUDp, SSp-bfd, SSp-ul, SSp-m, and SSp-n varied between 1.19 and 4.61 (Figure 1). Areas with the highest TD were ILA, ORBm, and PL, with values equal to 4.61, 4.01, and 3.57 respectively. Areas with the lowest TD were SSp-n, AId, and VISli, with values equal to 1.19, 1.21, and 1.28 respectively. Although TD provides a summary and a fair comparison of the dimensions of different cortical areas, significant variance was also observed in higher dimensions (Figure 1B).

**Figure 1:**
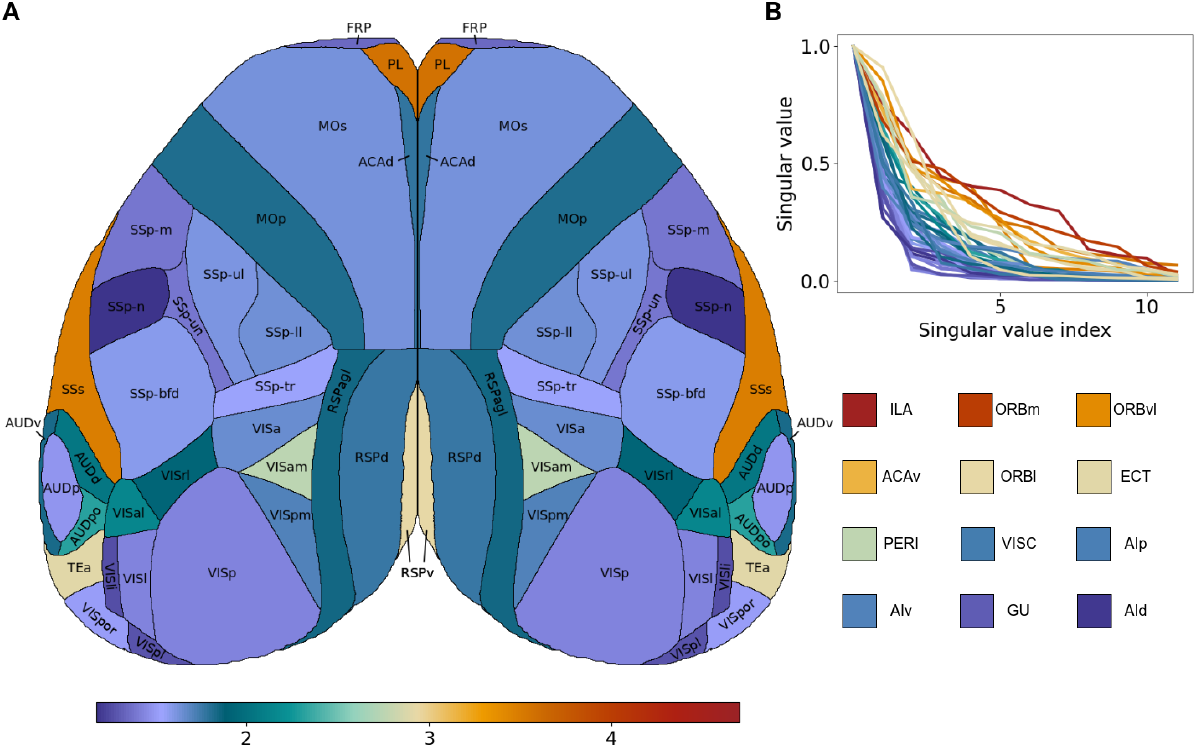
(A) TD of individual cortical areas with top-down view generated using [40]. Areas not visible from the top are shown as squares of the representative color at the bottom of the figure. (B) Normalized singular values of different cortical areas demonstrating the distribution of variance across dimensions.

The coordinates of inputs to high-dimensional areas varied with higher spatial frequency than those of inputs to low-dimensional areas (Supplement Figure 13).

We also calculated TD separately for each cortical layer within every cortical area. Layer-wise TDs had a similar range of values and were strongly correlated with area-wise TDs (Figure 2).

**Figure 2:**
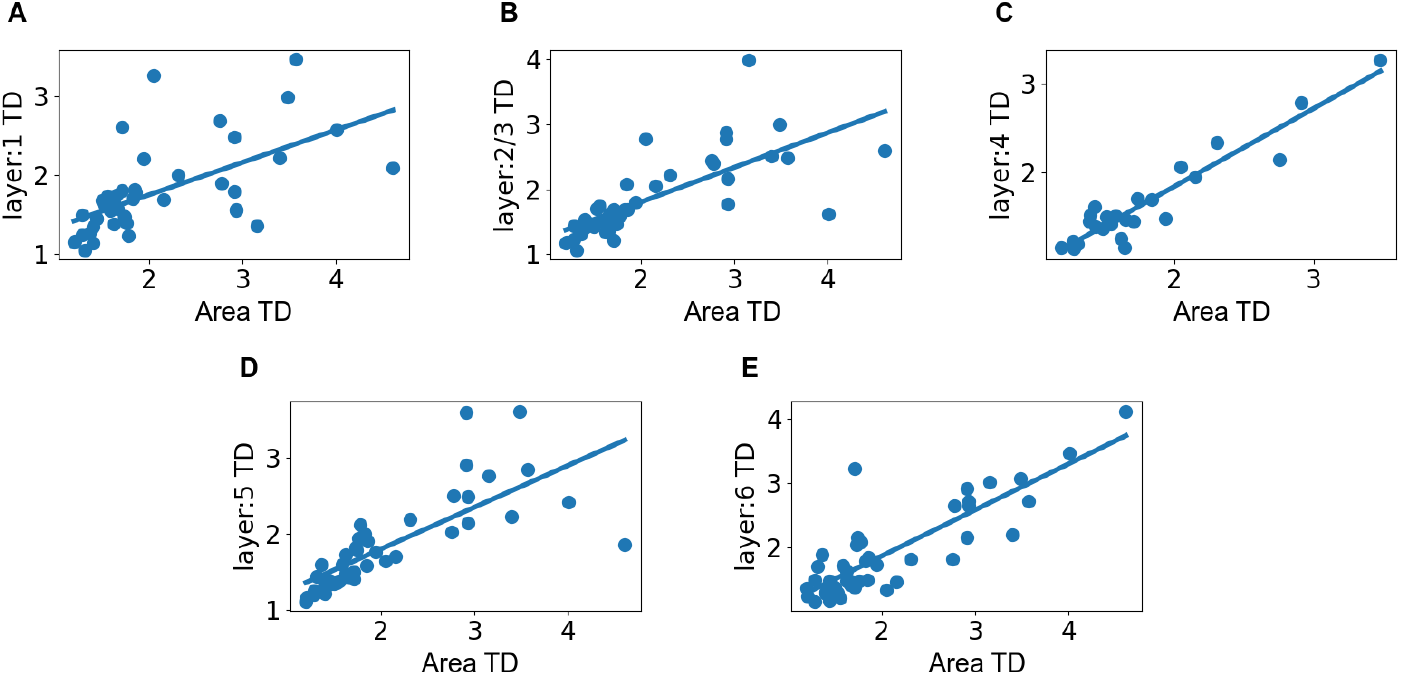
Layer-wise TD vs. area-wise TD with the line of best fit demonstrating positive slopes for all layers with 95% confidence interval of (A) [0.24, 0.58] for layer 1, (B) [0.37, 0.69] for layer 2/3, (C) [0.75, 1.02] for layer 4, (D) [0.4, 0.7] for layer 5, and (E) [0.57, 0.86] for layer 6. The range of area TD is smaller for Layer 4 because the highest-TD areas are agranular frontal areas.

The mesoscale connection model [30] is based on interpolation of finite tracer experiments from [44]. For example, these experiments include 33 injections into VISp but only three injections each to SSP-n and SSP-ul. Areas with small numbers of injections may cause underestimation of the dimension of projections from these areas, and the results may also be sensitive to the details of individual experiments. To characterize the sensitivity of TD estimates to the presence of individual injections, we recalculated the TD after removing from the model (one at a time) each of the three experiments in each primary sensory area that contributed most strongly to the connection weights. The resulting dimensions for different areas showed little variation, with standard deviations averaging 0.1 dimensions across cortical areas (range 0.015-0.34). The two areas with greatest standard deviations were VISa and ILA; other standard deviations were less than 0.28.

### 3.2 Column-wise averaging of sensory coordinates

TD reflects the convergence of multisensory projections onto 100×100*µ*m voxels. It may not reflect the convergence of information within neurons, due to the spread of dendritic arbors beyond individual voxels, and local connections that spread information within cortical columns. To estimate the potential impacts of these factors, we combined the coordinates of voxels in the same cortical column in a weighted average. Specifically, we recalculated the coordinates of each voxel as a weighted average of coordinates of surrounding voxels, using a Gaussian function of lateral distance as the weighting function (see Methods Section 5.3). A small drop in TD due to column averaging would suggest that much of the variation in sensory coordinates occurred between columns, while a large drop would suggest substantial variation in coordinates within a column, perhaps including the convergence of diverse coordinates onto single neurons. Column-averaging typically reduced TD by 0.46 ± 0.58 (mean ± standard deviation)(Supplement Figure 11), with the greatest reduction in prefrontal areas ILA, ORBm, and ORBvl, and the least reduction in VISpl, RSPd, and RSPagl, which are involved in vision and spatial cognition [19, 37, 60]. There was an intermediate impact in the ventral (granular) part of RSP, which is distinctly involved in contextual fear memory [64].

### 3.3 Relationship between TD and cortical hierarchy

We also observed that TD tended to be greater for areas higher in cortical hierarchy. Figure 3 plots TD vs. the hierarchy score (HS) from [21], which is based on laminar connection patterns. The line of best fit has a positive slope, with a 95% confidence interval of [0.4-2.93], indicating a clear positive correlation between TD and HS across cortical areas. This relationship was also significant within individual cortical layers 5 and 6 (Figure 3). The trends in other layers were also positive, but not significantly different from zero.

**Figure 3:**
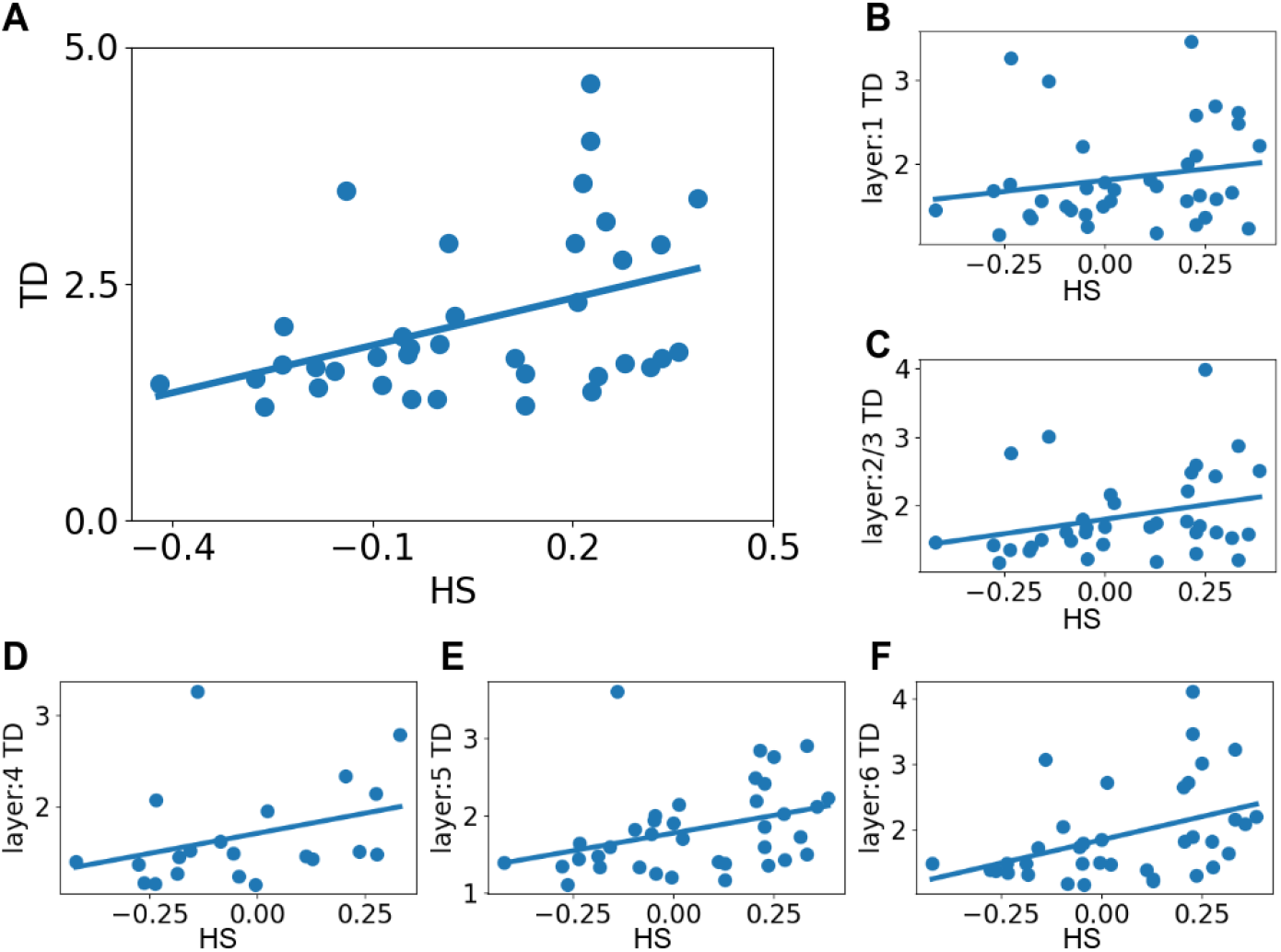
(A) Scatterplot and line of best fit of TD vs. hierarchy score (HS) for different cortical areas. The line of best has a positive slope with 95% confidence interval [0.4-2.93]. (B-F) Layer-wise TD vs. area HS. The lines of best fit have positive slopes for layers 5 and 6 with 95% confidence intervals of [0.07, 1.76], and [0.36, 2.49] respectively. The lines of best fit for layers 1, 2/3, and 4 were not significantly positive (95% confidence interval [-0.39, 1.47], [-0.1, 1.79], [-0.3, 2.07]).

### 3.4 Factors limiting topography dimension

In principle, TD values could have been as high as 12, as we analyzed 2D cortical coordinates of six primary areas. However, all TDs were substantially lower. This raises the question of which factors substantially limited TD, e.g., correlations between coordinates arising from different sources, or differences in connection strengths.

We decomposed the missing dimension of each area, 12-TD, into various factors, by making perturbations to the voxels’ 12D coordinates and weights and then recalculating the TD. We considered four factors: 1) Within-modality coordinate correlations; 2) Between-modality coordinate correlations; 3) Between-modality differences in mean connection weights; and 4) Within-modality differences in connection weights across voxels within a target area. The latter factor could affect TD if a connection were topographically organized with one part of the topography more strongly connected than another part, as in [38].

Figure 4 shows the TD of each area along with the missing dimension decomposed into these four factors. In many areas, the factor that most limited TD was non-uniform connection weights within modalities. This reduced TD by 3.4 +/-0.62 (mean +/-standard deviation). TD was also substantially limited by differences in connection weights between modalities (3.2 +/-0/84). Notably, linear correlations between coordinates of different modalities had relatively little impact on TD overall (1.5 +/-0.57), although they reduced TD by *>* 3 in the unassigned division of the primary somatosensory area.

**Figure 4:**
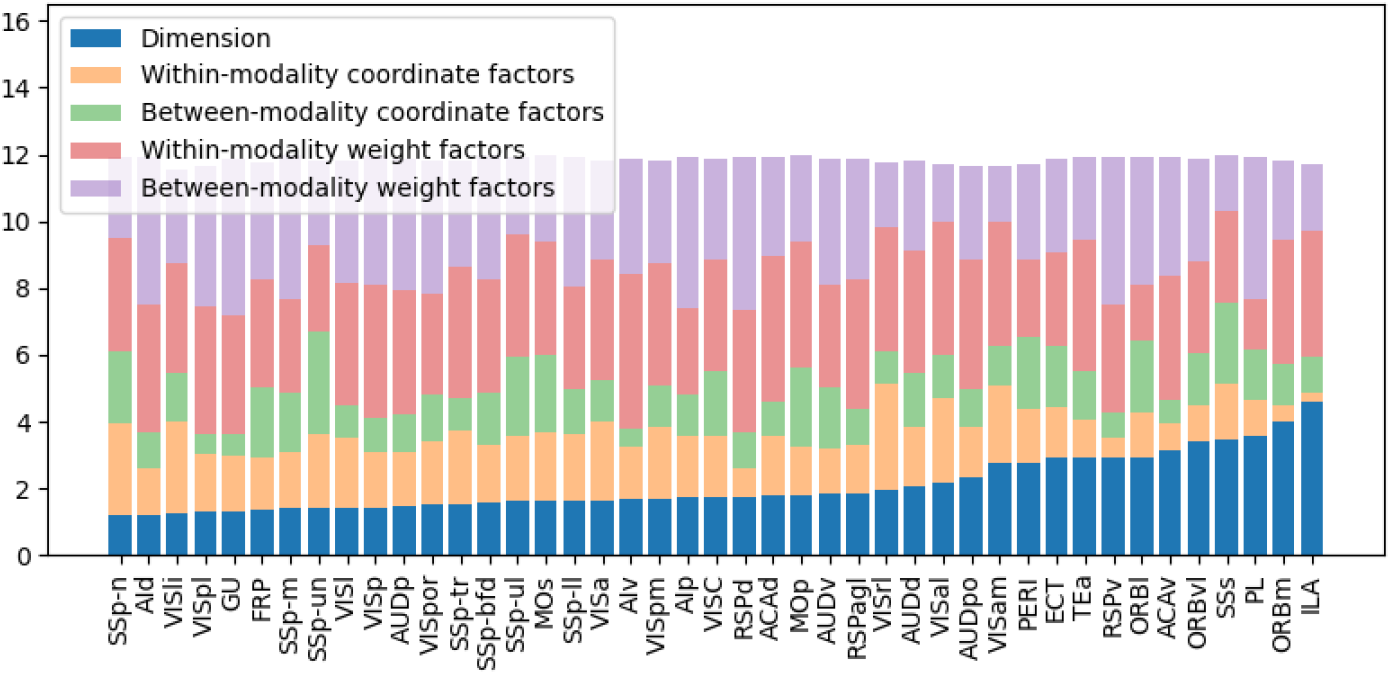
TD of each cortical area, stacked with missing dimensionality attributable to various factors, as described in the text.

### 3.5 Indirect connections

While direct connections from primary sensory cortical areas terminate extensively throughout the mouse cortex, information from these areas also spreads through indirect connections, which are sometimes stronger. As described in the Methods Section 5.3, we propagated primary sensory coordinates twice through the mesoscale connection model to study secondary connections. The topography of secondary connections exhibited strong correlations with the topography of primary connections. Figure 5 shows the distribution of primary vs. secondary connections from VISp. Coordinates of direct and indirect connections had strong positive correlations (*r*_*AP*_ = 0.827 and *r*_*ML*_ = 0.851 for anteroposterior and mediolateral coordinates, respectively). However, secondary coordinates had smaller ranges. Similar relationships were found between coordinates of direct and indirect connections from other primary sensory areas (Supplement Section 7.6).

**Figure 5:**
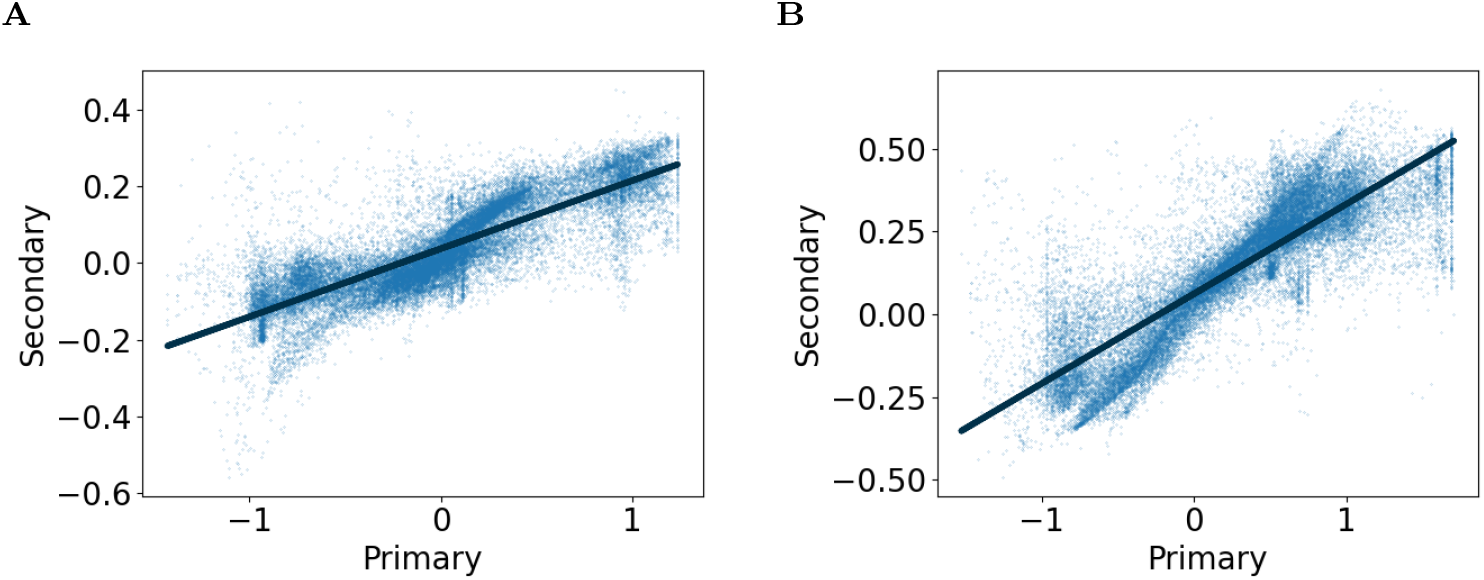
Primary vs. secondary propagated coordinates (normalized with unit variance) for VISp with the line of best fit demonstrating positive slopes with 95% CI of (A) [0.176, 0.179] for anteroposterior and (B) [0.269, 0.273] for mediolateral coordinates, and Pearson correlation coefficient of (A) 0.827 for anteroposterior and (B) 0.851 for mediolateral coordinates.

### 3.6 Topography dimension vs. neural activity dimension

We also sought the functional correlates of TD, which is a structural measure. One of the most fundamental properties of neural population activity is its dimension. This property can be meaningfully compared across the cortex, and different dimensions may optimally support different functions [16]. We suspected that higher TD might be associated with higher neural activity dimension (NAD).

Using Neuropixels recordings of spontaneous activity from [62], we constructed neural activity matrices from 100 neurons per area and 300 values of spike rate in consecutive non-overlapping 125ms windows similar to [51]. The eigenvalues of the covariance matrices of this activity were used to calculate the NAD using the participation ratio. The different numbers of neural recordings per cortical area result in different numbers of NAD values. To create balanced datasets, we randomly sampled an equal number of data points for each cortical area and repeated the sampling process 100 times. We performed linear regression of NAD values (derived from multiple neural activity matrices per cortical area) from TD for each of these 100 draws. We report the variance-weighted mean of the resulting regression coefficients. The line of best fit had a positive slope, with a 95% confidence interval of [1.0925, 1.0937], indicating a significant positive correlation between neural activity and topography dimensions. We repeated this analysis with window lengths of 250ms and 500ms and found significant positive slopes in each case. We then repeated the analysis using robust regression with Huber loss [23]. Again, the positive slope was significant with each choice of window length (Figure 6). Thus TD is significantly and robustly related to NAD.

**Figure 6:**
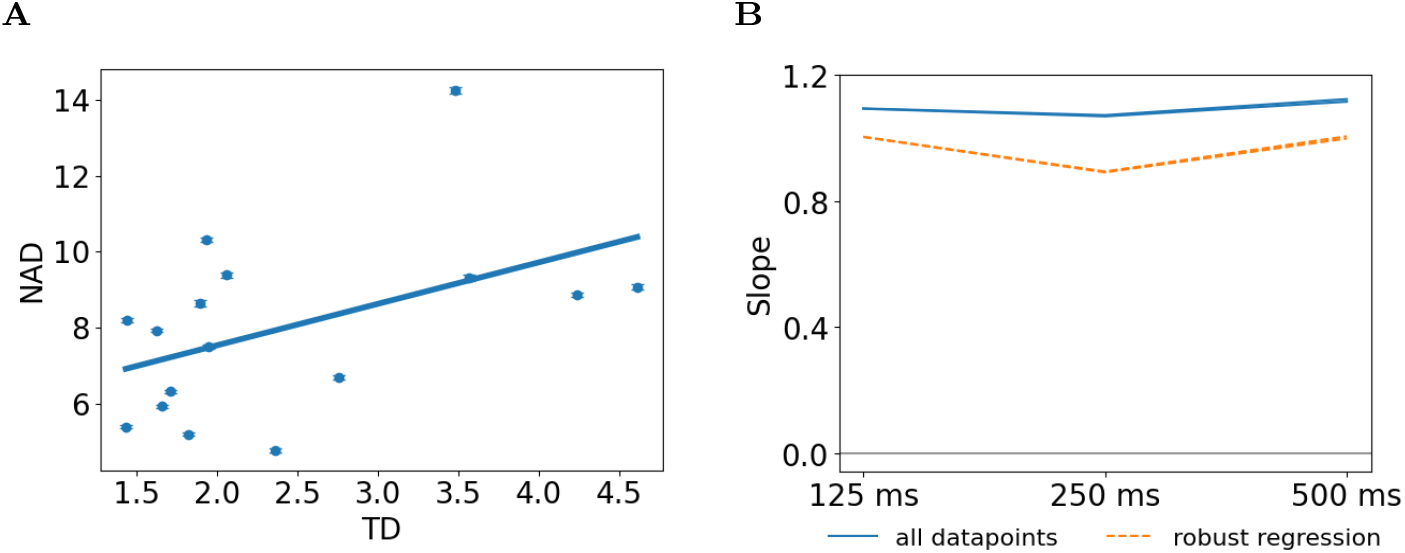
(A) Mean neural activity dimension (NAD) vs. TD for various cortical areas. The error bars indicate the mean +/-standard error over 100 neurons by 300 125ms time bin chunks of spontaneous spike-rate activity for each area. The line represents line of best fit for all points, and the shaded areas shows 95% confidence interval of [1.0925, 1.0937] (B) 95% confidence intervals of slopes for NAD vs. TD of all points (blue solid) and robust regression (orange dashed) for different window lengths. The gray line represents a slope of 0.

Because TD is correlated with hierarchy score (HS), we wondered whether the correlation between NAD and TD might be an artefact of a stronger cor-relation between NAD and HS. We therefore repeated the regression analyses with HS rather than TD as the regressor. NAD was also significantly related to HS for all window lengths, for both standard and robust regression (Supplement Section 7.7). However, the sign of the slope varied across analysis conditions.

Finally, we performed likelihood ratio tests to determine whether NAD was better predicted by both structural variables (TD and HS) together than by each one individually. These tests revealed that the full model was not significantly more predictive than the TD-only model, with p ∈ [0.2, 1] across window lengths and standard/robust regression (Supplementary Table 1). However, the full model was significantly more predictive than the HS-only model (p ∈ [0, 0.0066]).

#### 3.6.1 Population Sparseness

Another fundamental property of population activity is sparseness. Higher sparseness is associated with more efficient coding, and lower sparseness with greater representational capacity relative to population size [3].

Again using data from [62], we computed population sparseness, *S*^*P*^, of spontaneous activity using 125ms windows to estimate spike rates. We analysed two population-sparseness measures, 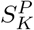 and 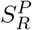, based on kurtosis and activity fraction, respectively (Methods Section 5.6.2). Figure 7A and 7C show scatterplots of *S*_*K*_ and *S*_*R*_ vs. TD in each cortical area. In each case the line of best fit for all data points was significantly negative (95% confidence intervals [-1.8530, -1.8529] and [-0.034, -0.034], respectively). The negative correlations persisted with window sizes of 250ms and 500ms, and with robust regression (Figure 7B and 7D). Thus, population sparseness is robustly related to TD.

**Figure 7:**
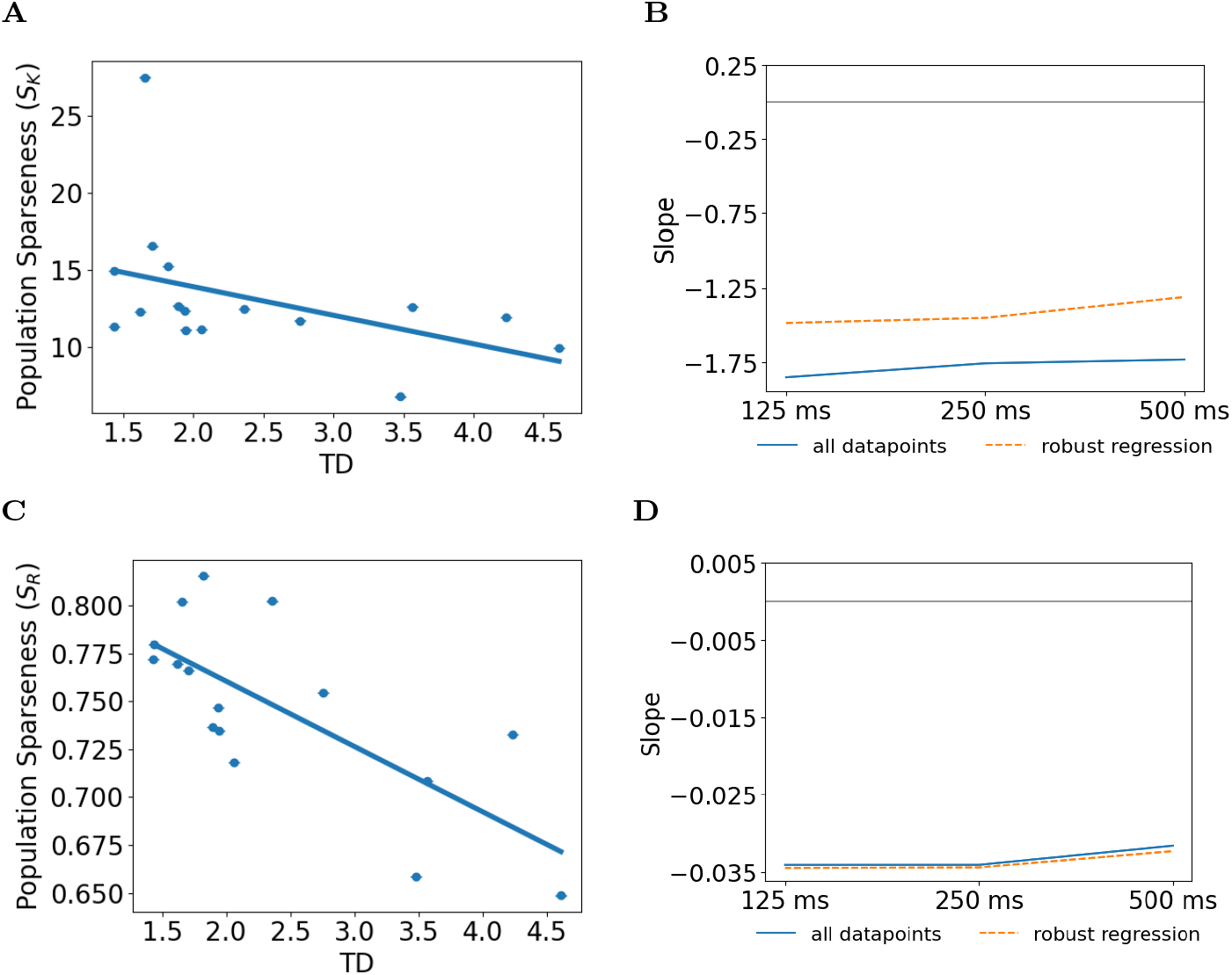
Population sparseness via (A) kurtosis, (C) activity fraction vs. TD. Sparseness measures were calculated using 125ms windows. The markers and error bars show mean +/-standard error of the mean for each cortical area. The lines represent line of best fit through all points for each area. The shaded regions illustrate 95% confidence intervals for the slopes, which are [-1.8530, -1.8529] for kurtosis and [-0.034, -0.034] for activity fraction. Panels (B) and (D) show confidence intervals for different window lengths with the kurtosis and activity fraction measures, respectively. Blue indicates confidence intervals with standard linear regression and orange indicates confidence intervals with robust regression. The gray lines represent a slope value of zero.

Because TD is correlated with the hierarchy score from [21], its relationship with population sparseness could simply reflect a stronger correlation between sparseness and hierarchy score. However, the relationship between population sparseness and HS was less consistent across measures. In particular, standard regression coefficients were positive for 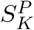 vs. HS for all window lengths, but coefficients were negative with 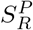 (Supplementary Table 4). Thus, the relationship between population sparseness and hierarchy score is less clear than the relationship between population sparseness and TD.

We performed likelihood ratio tests to determine whether both TD and HS together better predicted population sparseness measures than each measure alone. Both TD and HS contributed independently and significantly to prediction of population sparseness across all variations of window size, sparseness measure, and standard vs. robust regression (Supplementary Table 3).

#### 3.6.2 Lifetime Sparseness

We used the same neural activity matrices to estimate lifetime sparseness, *S*^*L*^. This is the sparseness of individual-neuron activity over time, averaged over neurons in each population. Figure 8A shows the relationship between mean lifetime sparseness, 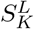 (calculated with 125ms window), and TD. The line of best fit had a negative slope with 95% confidence interval of [-0.4847, -0.4846]. The lifetime sparseness computed using activity fraction also exhibited a strong negative correlation with TD (Figure 8C). These negative correlations persisted with 250ms and 500ms windows and with robust regression (Figure 8B and 8D). Thus neurons in cortical areas with higher TD tended to have lower lifetime sparseness.

**Figure 8:**
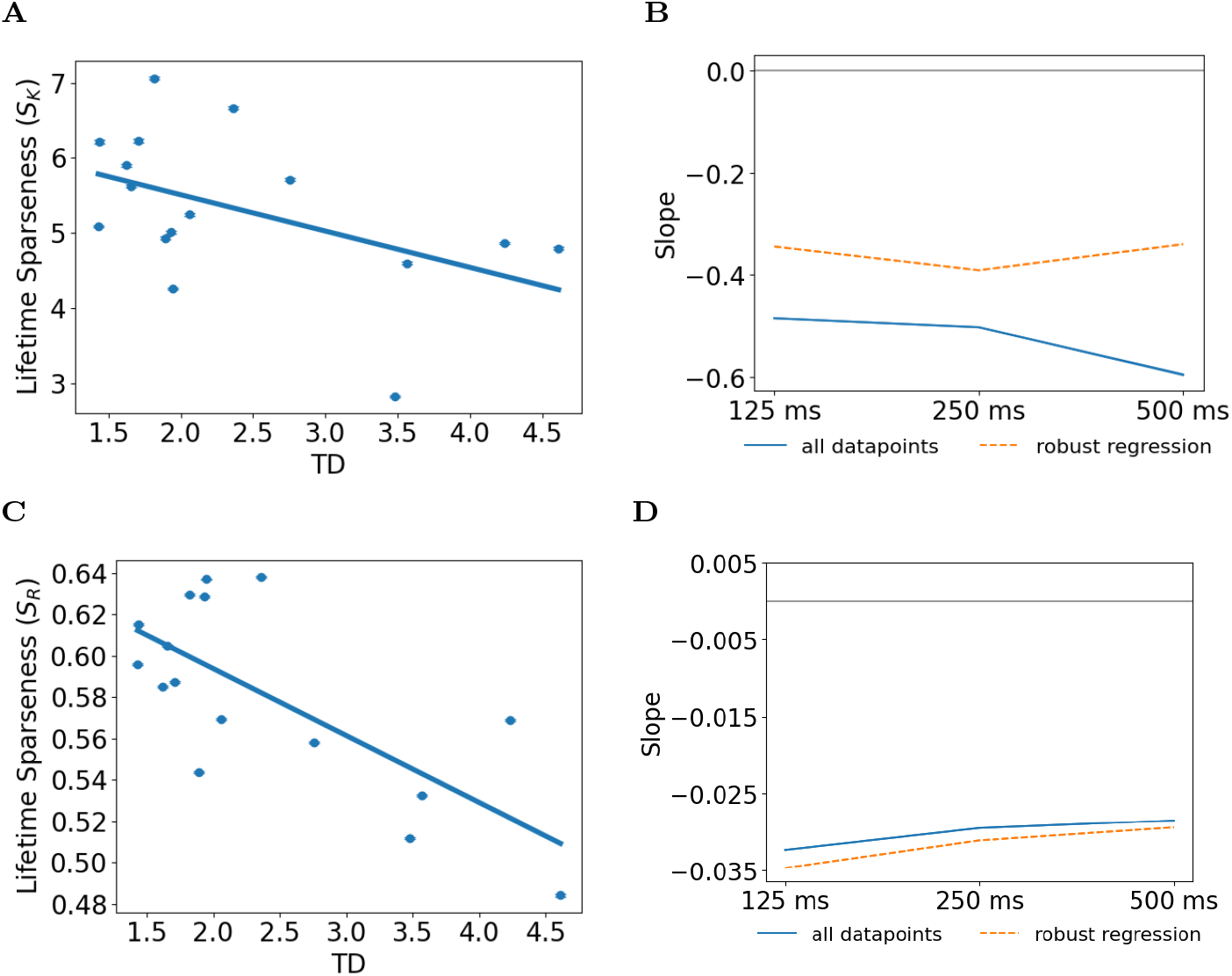
Lifetime sparseness (A) kurtosis, (C) activity fraction of mean points +/-standard error of the mean (spike rate measured in 125ms windows) vs TD. The lines represent lines of best fit for all points. The shaded regions illustrate the 95% confidence intervals of [-0.4847, -0.4846] for kurtosis and [-0.032, - 0.032] for activity fraction. Upper and lower ranges of 95% confidence intervals of the slopes of TD vs (B) kurtosis and (D) activity fraction of all points (blue solid) and robust regression (orange dashed) for window lengths. The gray lines represent the slope value of 0.

We also evaluated relationships between lifetime sparseness and HS and found that these were less consistent. In particular, the kurtosis measure of life-time sparseness, 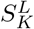, was positively correlated for standard regression against HS with window sizes 250ms and 500ms and robust regression against HS with window size 500ms, but negatively correlated otherwise (Supplement Table 6). The coefficient of the HS contribution to the full HS-TD model was also inconsistent across analysis conditions. In contrast, the coefficient of the TD model was consistently negative. TD contributed significantly in the full model, independently of HS, across sparseness scores, time windows, and standard/robust regression. HS contributed significantly with *p >* .05 in only 3/12 of these variations (Supplementary Table 5). Thus lifetime sparseness is robustly and significantly negatively correlated with TD, independent of HS, while its relationship with HS is less clear.

## 4 Discussion

Multiple organizational principles coexist in the anatomical mapping of connections. Using the strengths of connections between areas and graph theoretical measures, modules of organization can be defined [63]. Additionally, beyond the strength of connections, additional principles can be observed. For example, a hierarchical organization can be observed [12] even though feedforward and feedback connections between areas are of similar strengths, but different laminar patterns. Analysis of the mesoscopic connectome in the mouse cortex with an emphasis on laminar connection types leads to an unsupervised discovery of hierarchy [21]. In this study, we go beyond that. Connections between areas are not viewed just by their integrated strength or type, but by detailed patterns at much higher, voxel based resolution. This allows us to define new measures such as topography dimension (TD). While there is a correlation between hierarchy and TD, TD allows additional predictive power for functional measures including the neural activity dimension, population sparseness, and lifetime sparseness of neural activity.

TD tended to be higher for areas higher in the cortical hierarchy, perhaps reflecting differences in multisensory processing at different levels. One important multisensory process is cue integration, in which more reliable estimates of object or event properties are obtained by combining information across modalities. In this context, a key distinction is that integration may resemble maximum likelihood estimation [49], with estimates from different modalities combined according to their reliability, or multisensory causal inference [31], which additionally considers the probability that the signals arise from the same source. In humans, magnetoencephalogram data reflect both simple fusion and causal inference, with stronger reflections of causal inference over more anterior areas [5]. Further, it has been argued that different parts of cortex contribute processing of temporal, spatial, and semantic correspondences, and evidence accumulation, in addition to causal inference [43]. These distinctions focus on integration of redundant information. However, TD differences might also reflect convergence of multisensory signals for other purposes, such as estimates that combine non-redundant information (e.g. neck proprioception and vision to estimate locations in trunk coordinates), flexible modulation of decisions by specific multisensory contexts, or binding of non-redundant information into a unified percept [9]. Future work should investigate potential relationships between TD and different kinds of integration.

Much recent work has compared cortical circuits to deep networks. A key motivation is that deep networks currently provide the best accounts of cortical activity, even when they are not specifically optimized to do so, but rather optimized for task-related or self-supervised objectives [55, 36, 28, 6, 42]. Deep networks also show promise in modelling relationships between circuit properties and sophisticated, ethologically relevant functions [53, 27]. The structure of mouse visual cortex can be closely approximated by a convolutional network [58], but it is unclear whether convolutional networks can approximate multisensory influences throughout the cortex. In contrast with the correlation between TD and hierarchy score, the convolution dimension does not typically increase with depth in such networks. Also, convolution over multiple dimensions would involve combinatorial explosion of layer size. However, qualitatively, many of the coordinate maps we observed (e.g., Supplement Figure 12) resembled gradient images that have been randomly remapped with various spatial frequencies. Perhaps this multisensory architecture could be approximated by applying different image remappings to convolution results that converge from different modalities. Numerical experiments suggest that this model can produce high-dimensional topographies in which dimension varies reliably with spatial frequency content of the remapping functions (Supplement Section 7.10).

It is not just multiple sensory systems that are deeply intertwined, but motor, executive, memory, affective, and sensory systems together [44, 69]. Future analysis and modelling will have to address the principles and statistics of these broader interactions in a similar manner. Looking farther ahead, the spectrum of future computational models should include large models that perform realistically sophisticated behaviour, carefully grounded in anatomy and physiology in order to provide realistic explanations. In contrast, advanced deep networks such as transformers are increasingly capable of approximating brain function [71, 1], but their starkly non-biological structures make them inadequate brain models. Future work should expand the characterization of whole-brain connectivity beyond connection strength [44], and beyond sensory sources, to discover broader principles of connection structure that provide a better grounding.

As a next step in this direction, the present work should be extended to more thoroughly analyze indirect multisensory connections. We showed here that sensory coordinates arising from secondary connections are correlated with those of primary connections, but vary less widely. Coordinates due to tertiary connections (not shown) continue this trend, becoming less topographically organized. Future work should characterize these connections more thoroughly, and attempt to reconcile them with spatiotemporal receptive fields.

In contrast with HS, TD had clear and significant relationships with all of the fundamental functional measures we examined (neural activity dimension, population sparseness, and lifetime sparseness). Although TD is a simple and well motivated structural measure, future work may reveal alternative, perhaps related measures, which are yet more strongly predictive of these properties.

## 5 Methods

### 5.1 Mesoscale connectivity model

We studied multisensory convergence using the mesoscale connectivity model of [30]. This is a voxel-to-voxel map of connection density that is based on 428 anterograde tract tracing experiments from the Allen Mouse Brain Connectivity Atlas [44]. Based on these experiments, [30] estimated voxel-to-voxel connection weights using Nadaraya-Watson kernel regression. The voxels in this model are 100*µm* on each side. Due to the number of experiments being much smaller than the number of voxels in the map, the resulting estimated projections are expected to be smoother than the real projections.

### 5.2 Flatmaps of primary sensory cortical areas

Our first analysis step was to define a 2D coordinate frame for each primary sensory cortical area in the right hemisphere. Separately for each primary area, we projected the area’s voxels onto the surface of the sphere that provided the best least-squares fit of that area’s voxel positions. Each voxel was then assigned mediolateral and anteroposterior coordinates. A voxel’s mediolateral coordinate corresponded to its angle to the right from the mid-sagittal plane, and the anteroposterior coordinate corresponded to its angle forward from vertical. These coordinates were then normalized by subtracting the area-wise mean and dividing by the area-wise standard deviation. This approach produced an essentially uniform coordinate density per unit cortical area, because primary sensory areas of mouse cortex are smooth and approximately semi-spherical (in contrast with, for example, orbito-frontal areas).

We analysed connections from six primary sensory areas (VISp, AUDp, SSp-bfd, SSp-ul, SSp-n, and SSp-m). We defer the study of piriform cortex because it has weak spatial organization, and other parts of SSp because preliminary analysis suggested weaker connection topography.

### 5.3 Sensory coordinate propagation

We propagated the flatmap coordinates through the mesoscale connection model [30] to each voxel in the isocortex. Specifically, we used voxel-to-voxel connection strengths to estimate the weighted input coordinates to each voxel. We then found the weighted mean of each coordinate from each primary sensory area, producing 12D sensory coordinates for each voxel in isocortex.

While our analysis focused on these direct-connection sensory coordinates, we also examined indirect projections. To do so, instead of propagating flatmap-based coordinates from primary sensory areas through the mesoscale model, we propagated weighted-mean coordinates from all cortical areas. We estimated secondary-connection coordinates of each voxel in the cortex as weighted means, again using weights from the mesoscale connectivity model, but weighing direct-connection coordinates instead of flatmap-based coordinates.

This method may not accurately reflect the convergence of multisensory information onto cells in each voxel, because synapses may occur in layers other than those that contain cell bodies, and information may be mixed by dense local connections. For these reasons, we performed an additional step in which we combined the coordinates of nearby voxels in a weighted average. The weights in this case were a Gaussian function of lateral inter-voxel distances, with a standard deviation of 97*µ*m, the mean width of interlaminar connection densities estimated by [58]. Each lateral connection distance was calculated as the distance between a neighboring voxel and a streamline that connected the central voxel to the inner and outer cortical surfaces. Streamlines followed the path of steepest ascent through the cortex, based on a solution of Laplace’s equation with boundary conditions at the inner and outer voxels, as in [26].

### 5.4 Topography dimension

Taking the 12D coordinates of all the voxels in a given area and weighing them by connection strength, we formed a 12 × *N* matrix (where *N* is the number of voxels in a given area) that encapsulates the distribution of the coordinates within each cortical area. We computed the dimension of each cortical area using the participation ratio (PR) [14] of coordinate matrix after zero-centering,

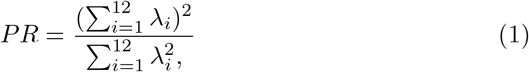

where *λ* are the eigenvalues of the covariance matrix. If 12D coordinates of different voxels in a cortical area were organized such that one eigenvalue could explain the variance of the covariance matrix, i.e. *λ*_*i*_ ≈ 0, for *i* = 2 … 12, the dimension of the cortical area would be close to 1. Alternatively, if all the eigenvalues contributed equally to the variance, i.e. *λ*_1_ = *λ*_2_ = … *λ*_12_, the dimension of the area would be 12. Otherwise, the dimension of the area would vary between 1 and 12.

Various other measures of dimension also have this property, and some have additional intuitive properties (Supplement Section 7.1). We report results based on the PR, which is conventional, but we experimented with two additional dimension measures and found broadly similar results, including patterns of correlation with neural activity data.

The PR value corresponds approximately to the number of dimensions required to explain 80% of population variance [14]. For the sake of brevity, we refer to the PR of an area’s coordinate matrix, derived from direct connections, as that area’s topography dimension (TD).

### 5.5 Decomposition of missing dimension

Because we analysed 2D coordinates projected from six primary cortical areas, TD could have been as high as 12, but in fact ranged from 1.2 to 4.6. We decomposed the missing dimension (12-TD) of each area into four factors by perturbing the coordinates and weights in various ways, as described below, and recalculating the TD.

#### Within-modality coordinate correlations

We whitened the coordinates within each modality using the SciPy stats package. This resulted in sensory coordinates with unit variance, and without within-modality correlations.

#### Between-modality coordinate correlations

To assess the impact of correlations between coordinates of different modalities, we shuffled the target voxels independently for each modality. Note that shuffling removes both linear and nonlinear correlations, but only linear correlations affect TD.

#### Within-modality weight imbalances

Dimension can be impacted by within-modality differences in connection weights across target voxels. To assess the impact of this factor, we normalized connection weights within modalities, so that each voxel in a target area received a connection of the same weight for a given modality. A minority of target voxels were originally unconnected to a given source, so that their coordinates for that source were undefined. After setting their weights to the mean, we therefore set their coordinates to random values with the same mean and variance as those of other target voxels in the area.

#### Between-modality weight imbalances

The calculation of TD involves scaling coordinates by connection weights, so that weaker connections contribute less to TD. Therefore, an area’s TD would be lower than the maximum if some modalities provided stronger input than others. To assess the impact of this factor, we divided connection weights by their modality-wise mean, so that connections from different modalities became evenly weighted.

Because some of these perturbations involved random variables, we report the mean of ten repeats with different random seeds. Standard deviations were generally less than 0.1 dimensions.

Applying all of these perturbations together brings an area’s TD to roughly 12. However, the difference in dimension due to a single perturbation may depend on co-occurrence with other perturbations. To assign a dimension to each perturbation, we first defined a 4D binary perturbation volume with various combinations of perturbations at the corners. We then averaged the difference with and without a given perturbation over all possible paths along the edges from zero to four perturbations. Using this approach, the sum of differences due to each perturbation equals the total effect of all perturbations. This is not generally true if one defines the effect of a given perturbation as the average of conditions with minus the average of conditions without.

### 5.6 Relationships between topography dimension and neural activity

We also examined the relationships between topography dimension and neural activity by analyzing Neuropixels data from [62]. This dataset contains sorted spikes from individual neurons. We focused on spontaneous activity before and between experimental trials. Recordings included 33 pre-trial periods with a mean duration of 501s (range 296-1016s), and 99 between-trial periods with a mean duration of 2213s (range 104-8840s). Trials included visual stimulation, choice presentation, and action, but we did not analyse trial data. This dataset is ideal for correlating with brain-wide TD because it contains extensive spontaneous activity and spans 16 cortical areas. The analysis focused on the dimension and sparseness of neural activity, fundamental measures that are relevant throughout the cortex.

#### 5.6.1 Neural activity dimension

We first organized the data into multiple matrices of neural activity for each cortical area. The matrix elements were spike rates (spikes per second) calculated over 125ms windows. Rows corresponded to different neurons, and columns to different non-overlapping contiguous time blocks spanning *T* (*s*). We experimented with different *T* ranging from 37.5s to 62.5s resulting in 300 to 500 spike rate samples respectively. Using shorter blocks allowed more trials (lower-variance estimate) but longer blocks could have revealed higher-dimensional activity (lower-bias estimate). We report results with windows of 125ms and blocks of 300 spike rate samples and show in the Supplement (Section 7.8) how estimated dimensions depend on both window length and *T*. The number of blocks differed between areas because not all areas were recorded in all experiments. Different numbers of neurons were recorded in different areas, ranging from 6 to 715. We randomly selected 100 neurons per cortical area, omitting areas with fewer neurons, to allow a fair comparison of dimensions across areas. We calculated the dimension of each block as the participation ratio of the corresponding matrix, leading to multiple estimates of neural activity dimension per cortical area. We repeated this process with 100 different random draws from the data, as described in Section 3.6. In order to combine the results of linear regression of NAD values from TD for each of these 100 draws, we used the variance-weighted mean to calculate the regression coefficient as,

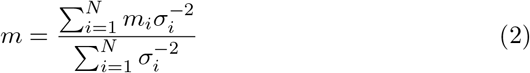

where *m*_*i*_ is the slope and 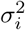 is the variance of uncertainty in the slope derived from the linear regression model for *i*^*th*^ draw. The 95% CI was computed as [*m* − 1.96*σ, m* + 1.96*σ*], where variance *σ* was calculated by 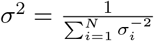.

#### 5.6.2 Neural activity sparseness

As there is no consensus measure of neural activity sparseness, we report two common measures, kurtosis and activity fraction. The kurtosis measure [48] is based on the shape of the distribution and is independent of its mean and variance. Given a list of *n* spike rates *r*_*i*_, it is defined as,

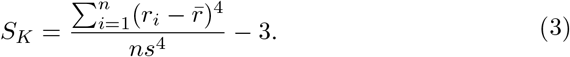

This is more specifically the excess kurtosis, beyond the kurtosis of a Gaussian distribution, which equals three. Distributions with large peaks and heavy tails have high kurtosis (high sparseness), while flatter distributions with thin tails have low kurtosis (low sparseness).

The activity fraction [66] is a rescaled version of [54] and is defined by,

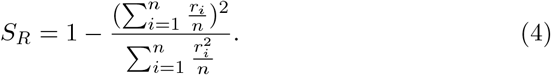

Flatter distributions have low *S*_*R*_, indicating lower sparsity. A high value of *S*_*R*_ occurs when the distribution contains rare values, suggesting higher sparsity.

We analysed both population sparseness and lifetime sparseness. Population sparseness describes instantaneous distributions of spike rates across a population of neurons (i.e., *r*_*i*_ correspond to different neurons). We report population sparseness of each block of data from each cortical area, averaged over time. Lifetime sparseness describes distributions of spike rates over time in a single neuron (i.e., *r*_*i*_ are spike rates of the same neuron in different time windows). We report lifetime sparseness averaged across neurons in each data block.

## 7 Supplementary information

### 7.1 Dimension measures

We defined TD as the participation ratio (PR) [14] of the 12 × *N* weighted coordinate matrix. The PR is calculated by first zero-centering and finding the correlation matrix and its eigenvalues, *λ*. Then,

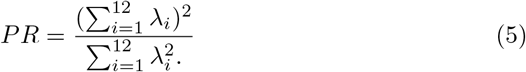

This dimension measure has been used repeatedly in neuroscience and it has some intuitive properties. In particular,

- If the variance is due to a single eigenvalue then the dimension is one,
- If the variance is due equally to all eigenvalues, the dimension is maximal (the length of the coordinate vectors),
- Otherwise it is between those values, becoming closer to the maximum as the the variance is more evenly distributed among different directions.

The PR is not unique in having these properties. Furthermore, the PR has a non-intuitive property. To illustrate, consider a two-dimensional distribution with a given standard deviation *σ* in one dimension and a potentially smaller standard deviation *ασ* in the other, where *α* ∈ [0, 1]. An intuitive dimension for such coordinates would be 1+*α*. However, the PR is less than this value for small *α* and greater for larger *α*. An alternative measure that has has dimension 1 + *α* asymptotically (and analogous properties in higher dimensions), while sharing the PR’s intuitive properties is,

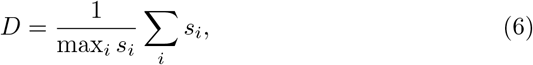

where *s* are singular values of the coordinate matrix.

Both of these measures are sensitive to uncertainty in the coordinates. To continue the motivating example, the sum-over-max measure is particularly sensitive if *α* is large. For example, suppose the true value of *α* is 1 but there is some error in the coordinates. Then the spread of the noisy data is likely to be empirically greater along one axis or the other, leading to estimated dimension *<* 2. This bias might be reduced given an estimate of the coordinate uncertainty.

To illustrate these differences, Figure 9 shows PR and sum-over-max measures for 1439 random coordinates drawn from a 12D Gaussian distribution. 1439 is the mean number of voxels in mouse isocortical areas in this study. The covariance matrix is diagonal with values [1, *α*^2^, …, *α*^2^]. Note that this is only a rough analogy to sensory coordinate data because in this illustration, error in the variance estimates arises from random sampling, whereas in the mouse data the entire population of voxels in each area is used but there is uncertainty in each coordinate.

**Figure 9:**
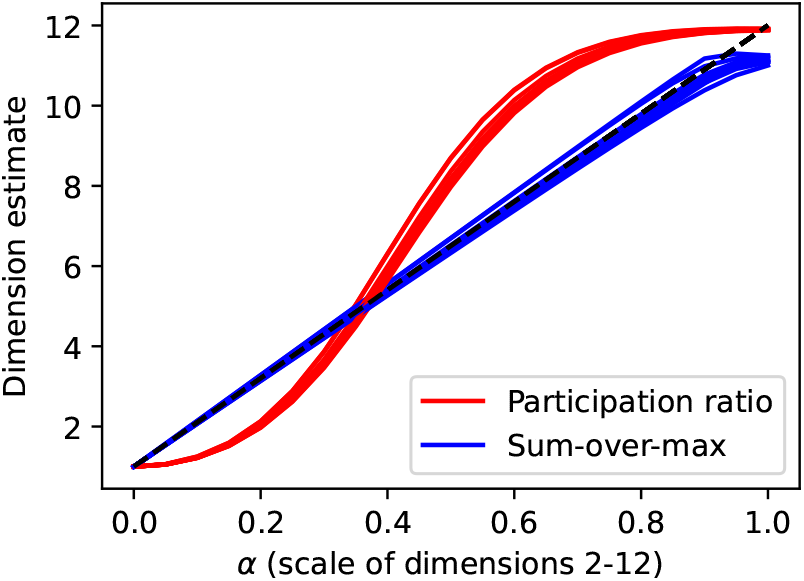
Illustration of different behavior of the PR and sum-over-max dimension measures with random Gaussian data. Random coordinate matrices are 12 × 1439, which is the average size of coordinate matrices in this study. Coordinates have standard deviation *σ* along the first axis and *ασ* along other axes. An intuitive dimension for such data would be 1 + 11*α*. The PR is systematically lower for small *α* and higher for large *α*. Sum-over-max follows 1 + 11*α* more closely, but underestimates it as *α* approaches 1. Each line shows dimension estimates for a given random draw of coordinates but varying *α* in steps of 0.05. Ten random draws are shown as different lines.

We experimented with both of these dimension measures. We also experimented with a modified PR that used singular values of the coordinate matrix rather than eigenvalues of the correlation matrix (this has the same intuitive properties as PR but deviates even further from the intuition of sum-over-max). The patterns of correlation with HS and with neural activity dimension and sparseness were broadly consistent across measures. We report results for PR as it is the most conventional measure.

### 7.2 Topography dimension

Figure 10 is a grayscale version of Figure 1 showing the TD of all cortical areas and normalized singular values of different cortical areas.

**Figure 10:**
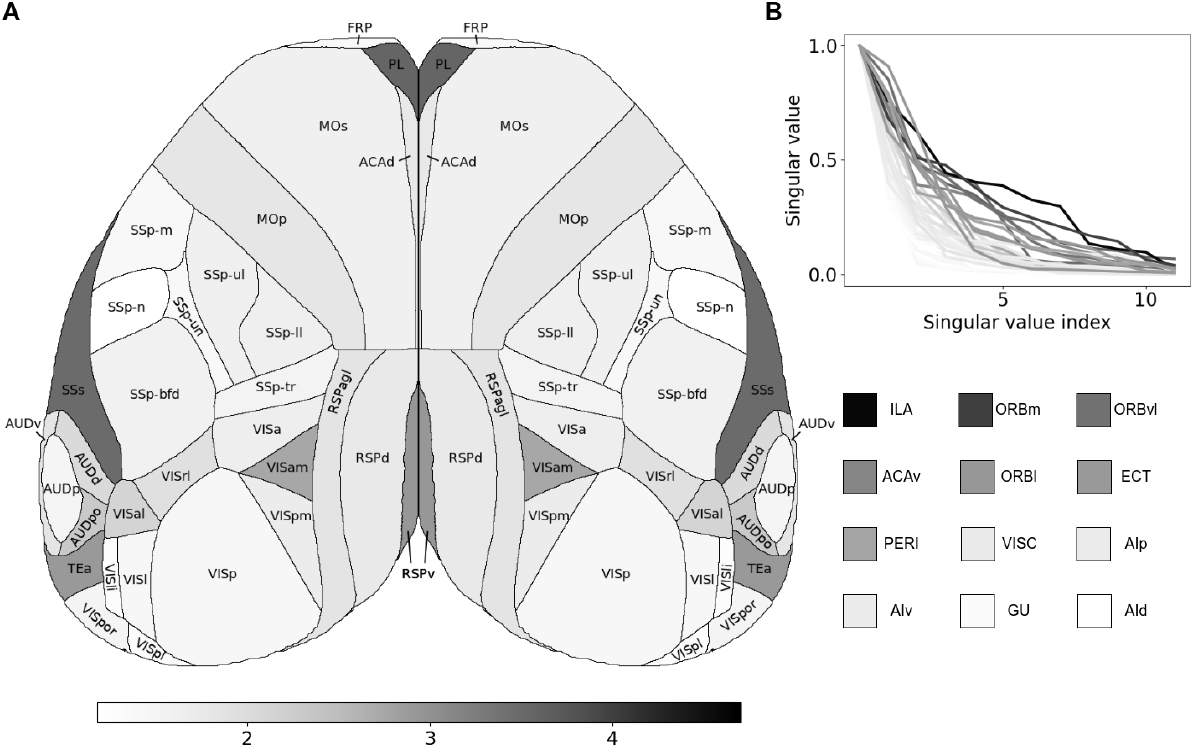
(A) TD of individual cortical areas with top-down view generated using [40] in grayscale. Areas not visible from the top are shown as square of the representative shade at the bottom of the figure. (B) Normalized singular values of different cortical areas demonstrating the distribution of variance across dimensions.

**Figure 11:**
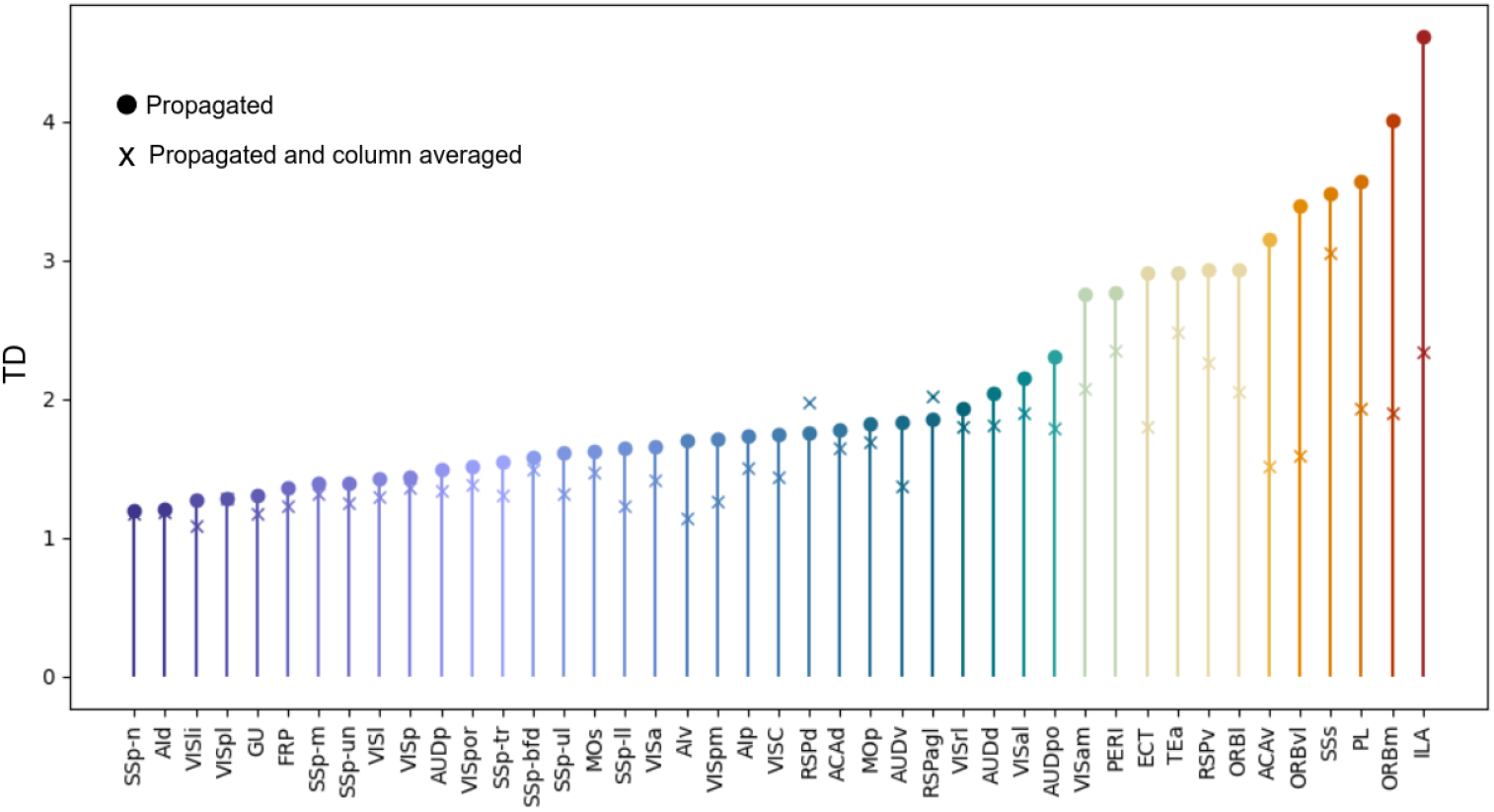
Dimension comparison between coordinates with and without averaging over cortical columns.

**Figure 12:**
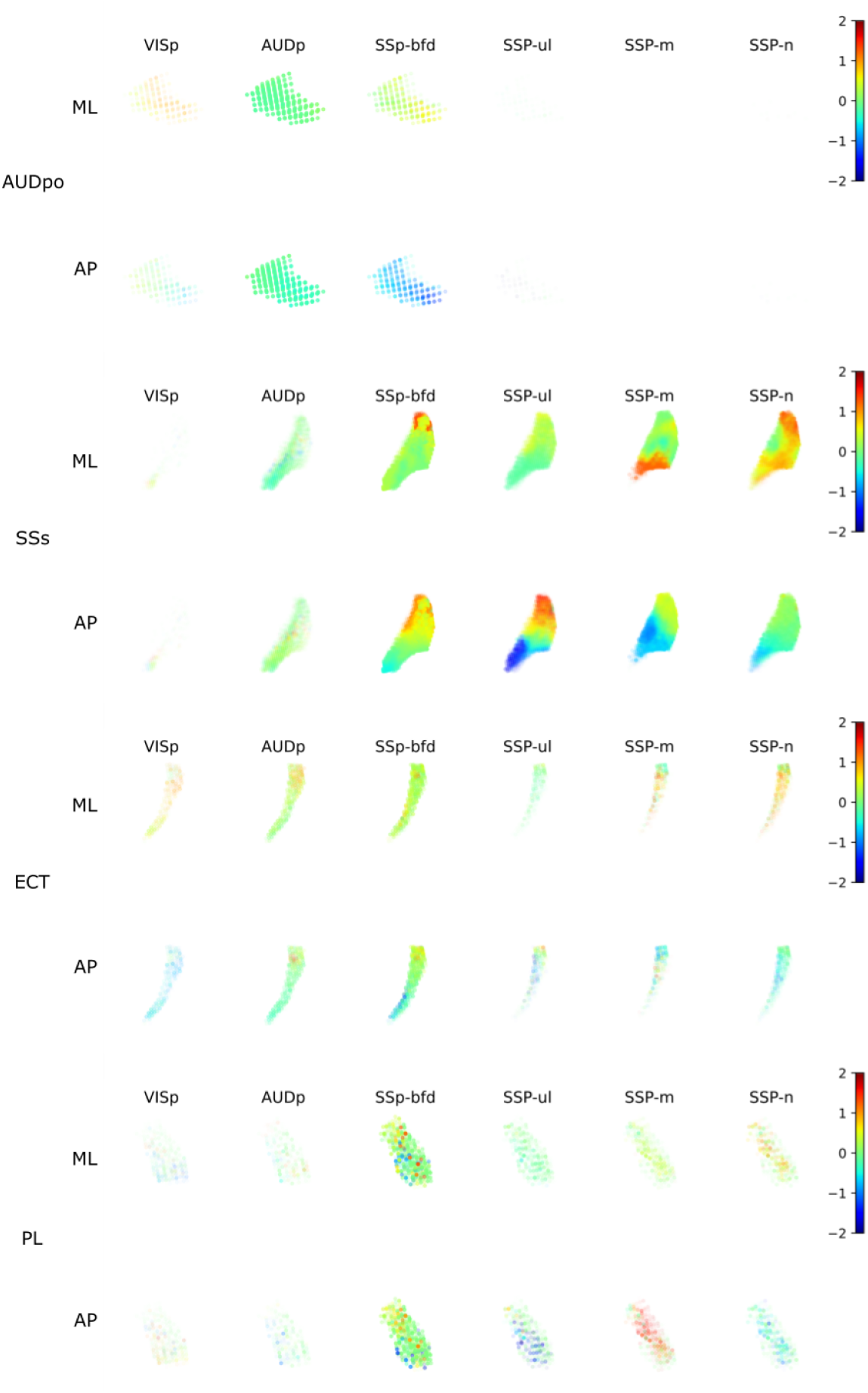
Primary sensory coordinates in four diverse cortical areas. From top to bottom, in order of increasing TD: posterior auditory area, secondary somatosensory area, ectorhinal area, and prelimbic area. Colored dots indicate normalized primary sensory coordinates of individual voxels in each of these areas. The 12 coordinates of each area are shown in a 2×6 grid with mediolateral coordinates in the top row, anteroposterior coordinates in the bottom row, and coordinates of projections from different primary sensory sensory areas in different columns, as labelled. The opacity of each point indicates strength of connection, normalized by the strongest connection into that area. These are dorsal views of right cortical areas from slightly behind and to the left.

**Figure 13:**
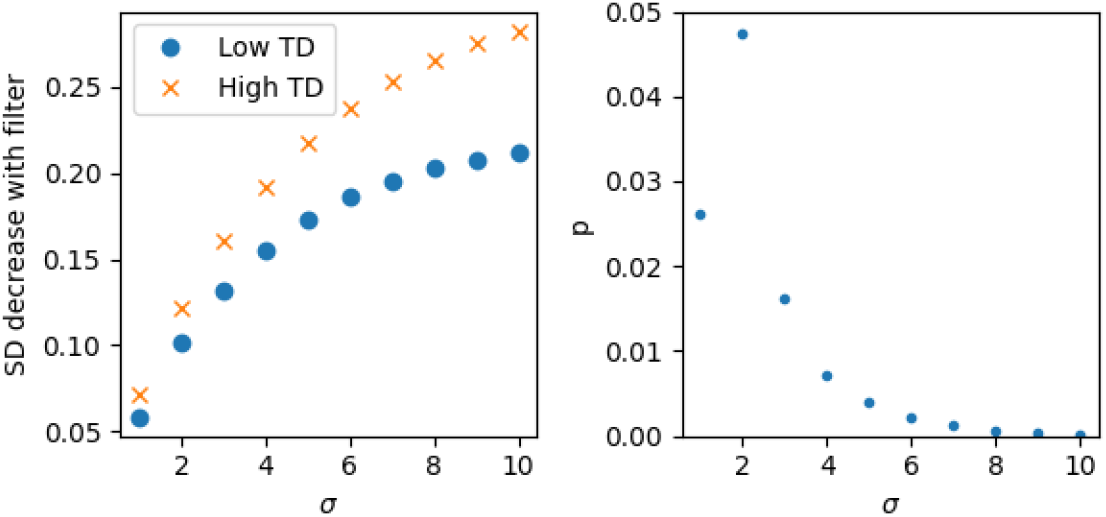
Coordinates of areas with higher TD tended to have higher spatial frequency content. We applied Gaussian filters with various *σ*, from 1-10 voxels, to coordinates in each area and assessed the effect on standard deviation of the coordinates. We calculated differences between coordinate standard deviations with and without the filter. The left panel shows the mean decrease in SD for low-TD and high-TD areas, for various filter *σ*. The right panel shows the probability that these decreases come from the same distribution, via two-tailed t-tests. All differences were significant with *p <* .05. We did not correct for multiple comparisons, because this was a single comparison tested for sensitivity to *σ*.

### 7.3 Column-averaged topography dimension

Figure 11 shows the TD of all cortical areas (circles) and corresponding TD after column averaging (x’s).

### 7.4 Plots of sensory coordinates

Figure 12 shows all 12 coordinates for each of four cortical areas.

### 7.5 Frequency content of sensory coordinates

Qualitatively, some areas with high TD appeared to have coordinates that varied with high spatial frequency. Area PL in Figure 12 is an example. To test this tendency, we applied spatial low-pass filters to the coordinates in each area, and quantified the resulting reduction in variance of the coordinates. Low-pass filtering more greatly reduces the magnitudes of higher-frequency coordinates. We used symmetric 3D Gaussian filters of various widths. We also accounted for the strengths of projections. In particular, we used Gaussian functions weighted by the connection weight, *w*, associated with each coordinate. Specifically, a filtered coordinate was,

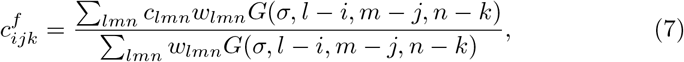

where *c*_*lmn*_ are the unfiltered coordinates of surrounding voxels, and *G* is the Gaussian function. We applied this filter independently to all 12 coordinates for each area. We then calculated the weighted standard deviation,

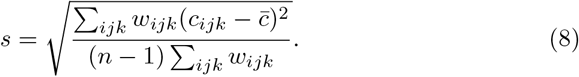

We calculated these standard deviations with and without the filter, and calculated their differences, 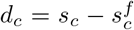, where *c* ∈ 1 … 12 is the coordinate index. We grouped together the *d*_*c*_ for the ten areas with lowest TD and ten areas with highest TD, yielding 120 differences per group. We then performed a two-tailed t-test to determine whether the filter more greatly attenuated coordinates (indicating higher frequency content) in the low-TD or high-TD group. For *σ* from 1 to 10 voxels (0.1 to 1mm), the high-TD group had significantly greater high-frequency content (Figure 13).

### 7.6 Relationships between coordinates of direct and indirect connections

The topography of secondary (indirect) connections is strongly correlated with the topography of primary connections. Scatterplots of primary vs. secondary visual coordinates are shown in Figure 5, and those of remaining modalities are shown in Figures 14 to 18.

**Figure 14:**
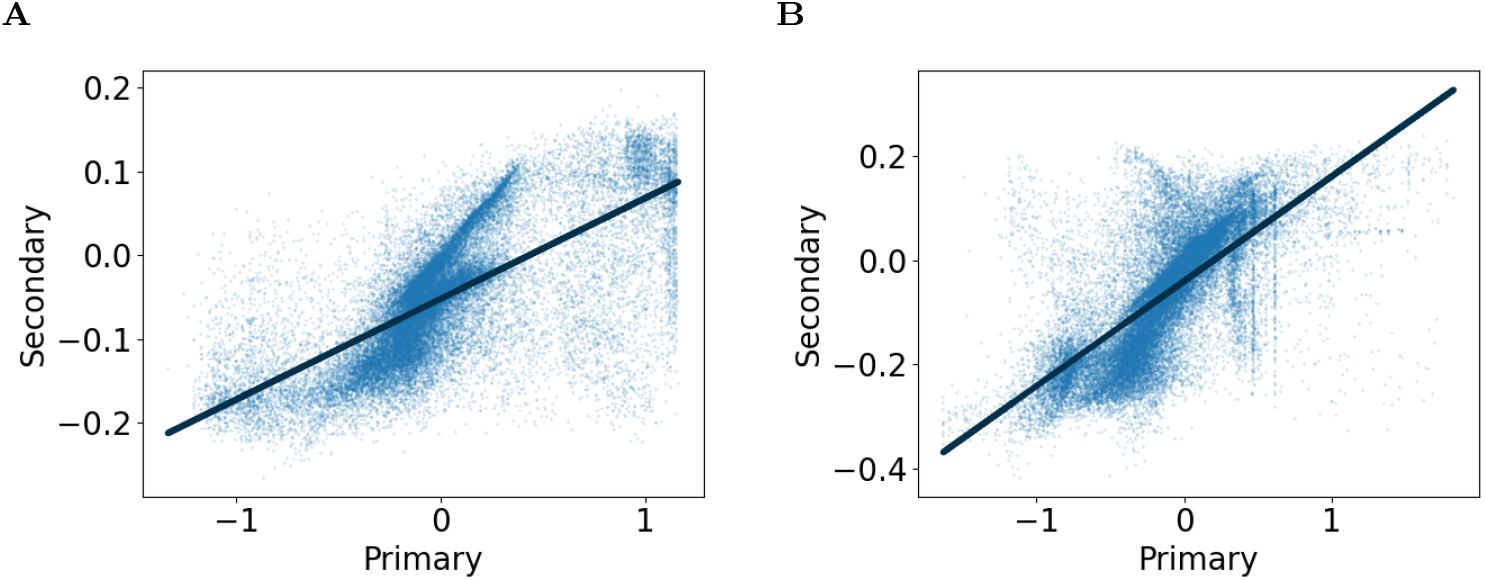
Primary vs. secondary propagated coordinates for AUDp with the line of best fit demonstrating positive slopes with 95% CI of (A) [0.119, 0.122] for anteroposterior and (B) [0.199, 0.203] for mediolateral coordinates, and Pearson correlation coefficient of (A) 0.691 for anteroposterior and (B) 0.698 for mediolateral coordinates.

**Figure 15:**
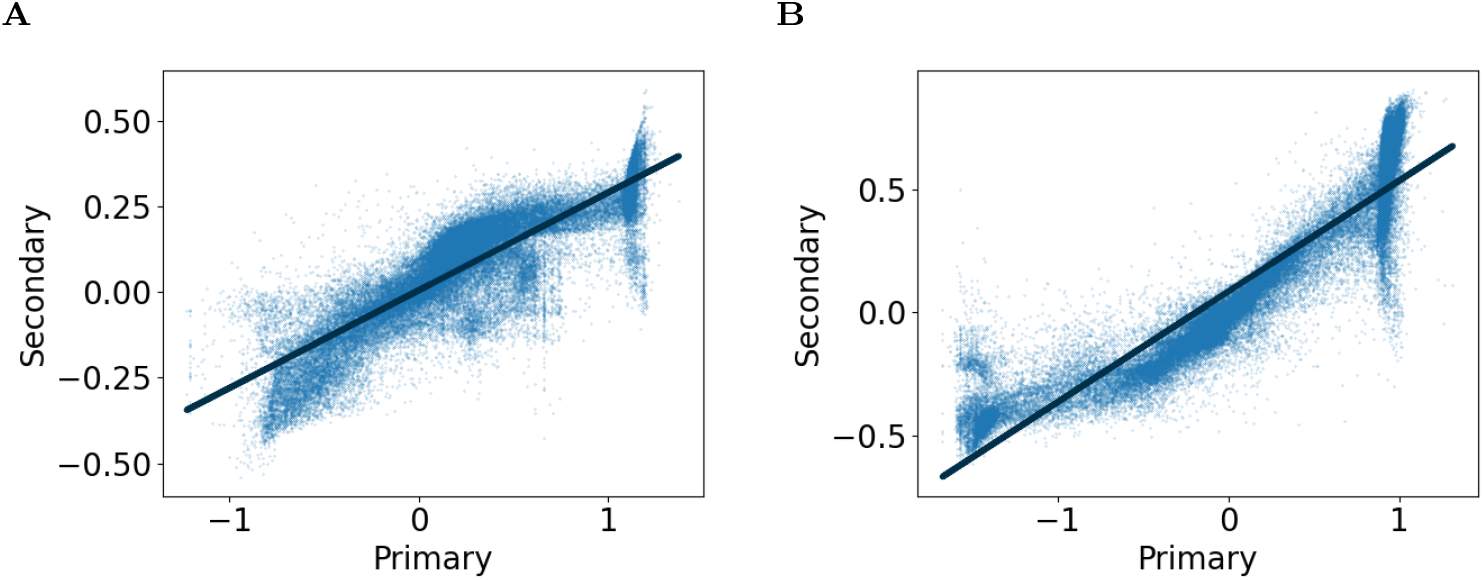
Primary vs. secondary propagated coordinates for SSp-bfd with the line of best fit demonstrating positive slopes with 95% CI of (A) [0.284, 0.287 for anteroposterior and (B) [0.447, 0.450] for mediolateral coordinates, and Pearson correlation coefficient of (A) 0.859 for anteroposterior and (B) 0.919 for mediolateral coordinates.

**Figure 16:**
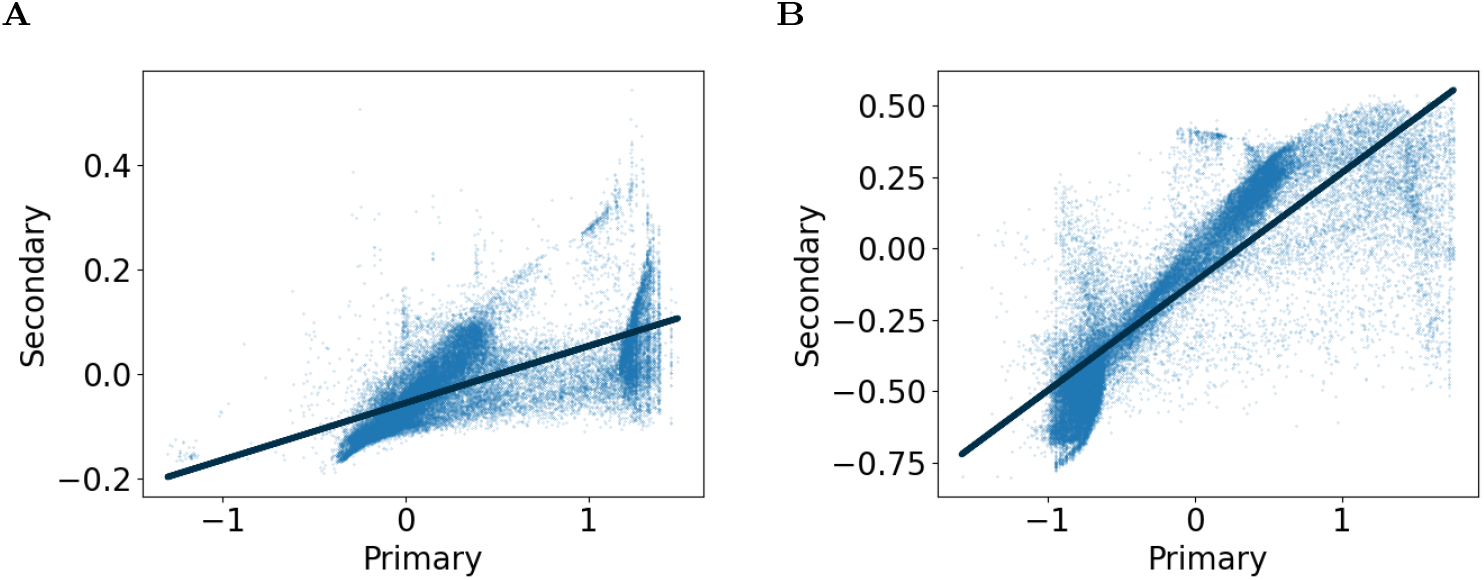
Primary vs. secondary propagated coordinates for SSp-m with the line of best fit demonstrating positive slopes with 95% CI of (A) [0.108, 0.110] for anteroposterior and (B) [0.380, 0.385] for mediolateral coordinates, and Pearson correlation coefficient of (A) 0.671 for anteroposterior and (B) 0.837 for mediolateral coordinates.

**Figure 17:**
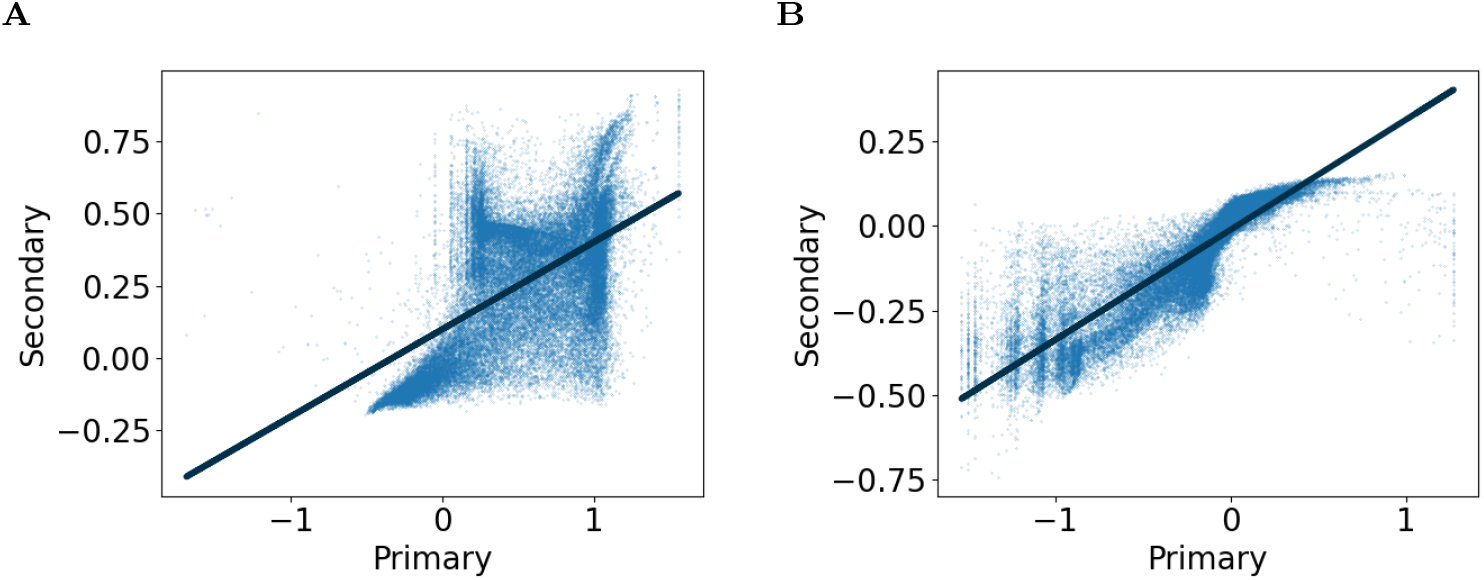
Primary vs. secondary propagated coordinates for SSp-n with the line of best fit demonstrating positive slopes with 95% CI of (A) [0.298, 0.306] for anteroposterior and (B) [0.325, 0.328] for mediolateral coordinates, and Pearson correlation coefficient of (A) 0.554 for anteroposterior and (B) 0.883 for mediolateral coordinates.

**Figure 18:**
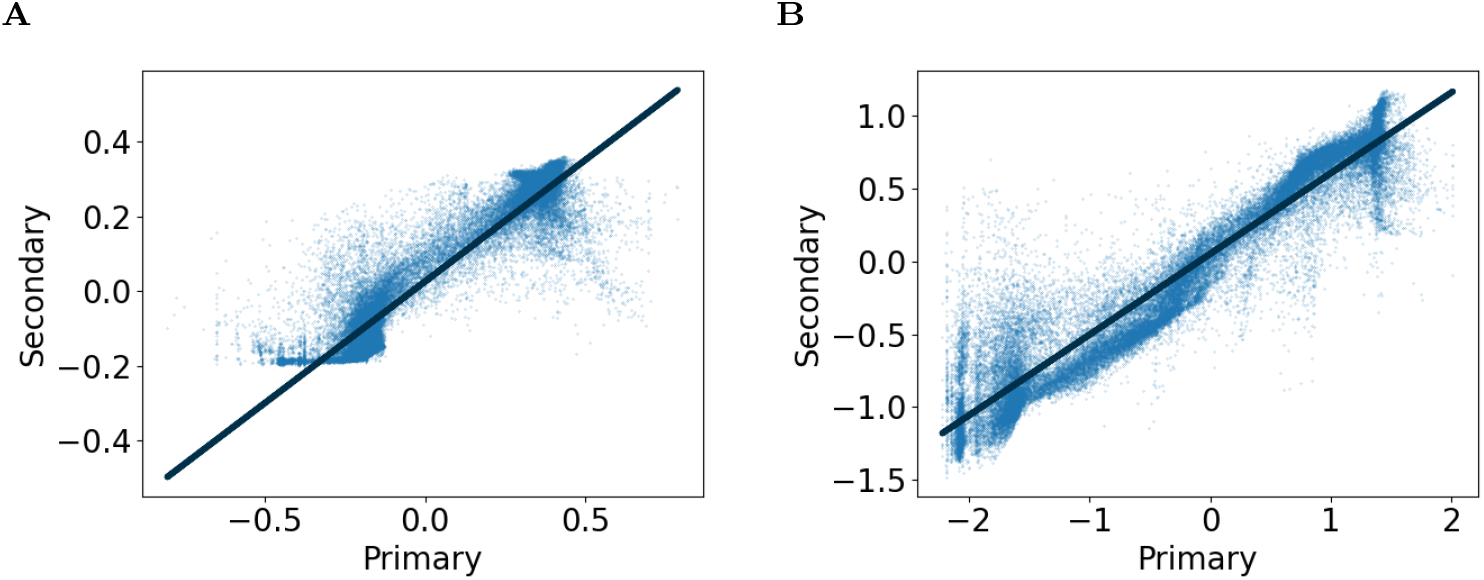
Primary vs. secondary propagated coordinates for SSp-ul with the line of best fit demonstrating positive slopes with 95% CI of (A) [0.650, 0.654] for anteroposterior and (B) [0.554, 0.557] for mediolateral coordinates, and Pearson correlation coefficient of (A) 0.927 for anteroposterior and (B) 0.950 for mediolateral coordinates.

### 7.7 Neural activity dimension vs. hierarchy score

The relationship between NAD and HS is characterized across different time windows in Figure 19 for both linear and robust regression.

**Figure 19:**
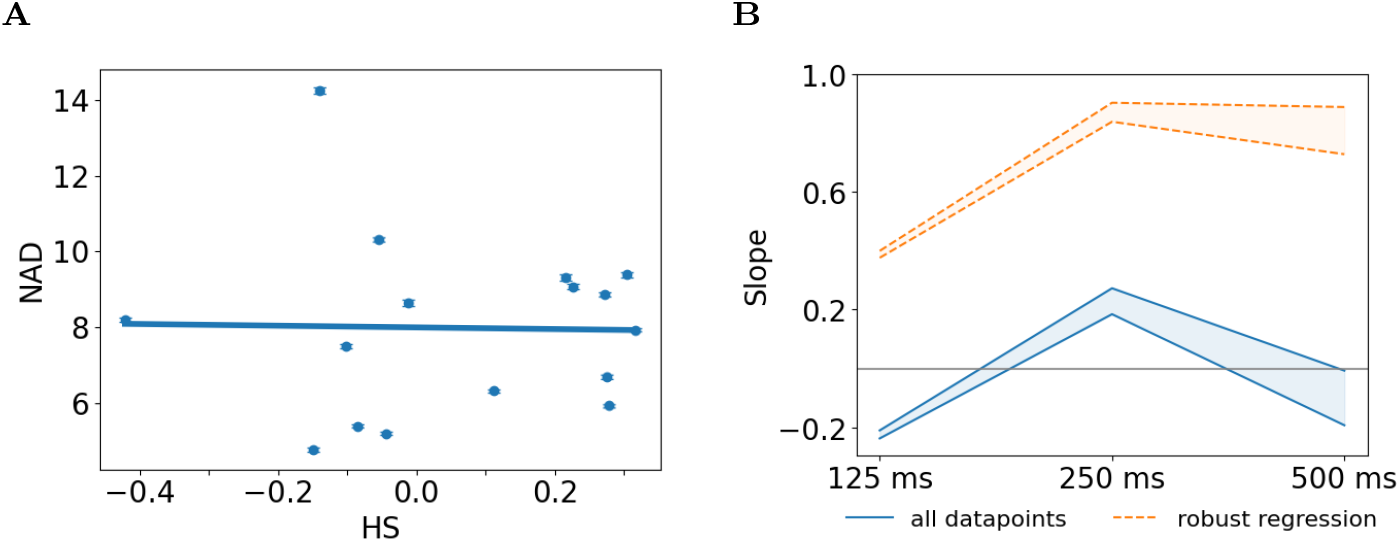
(A) Mean neural activity dimension (NAD) vs. HS for various cortical areas. The error bars indicate the mean +/-standard error over 100-neuron by 300-time-bin chunks of spontaneous spike-rate activity for each area, with 125ms bins. The line represents line of best fit for all points, and the shaded areas show 95% confidence interval of [-0.2377, -0.2105] (B) 95% confidence intervals of slopes for NAD vs. HS of all points (blue solid) and robust regression (orange dashed) for window lengths. The gray line represents a slope of 0.

### 7.8 Effect of number of spike rate samples on NAD vs. TD

In the main text we constructed spike-rate matrices with 100 neurons and 300 consecutive spike rates per neuron, with spike rates calculated over non-overlapping windows of 125ms to 500ms. To test sensitivity of the results to the numbers of spike rates in these matrices, we also constructed neural activity matrices from 100 neurons per area and 400 and 500 values of spike rate (again with spike rates calculated in consecutive non-overlapping 125ms to 500ms windows). NAD and TD remained significantly positively correlated in each of these variations, with both linear and robust regression (Figure 20).

**Figure 20:**
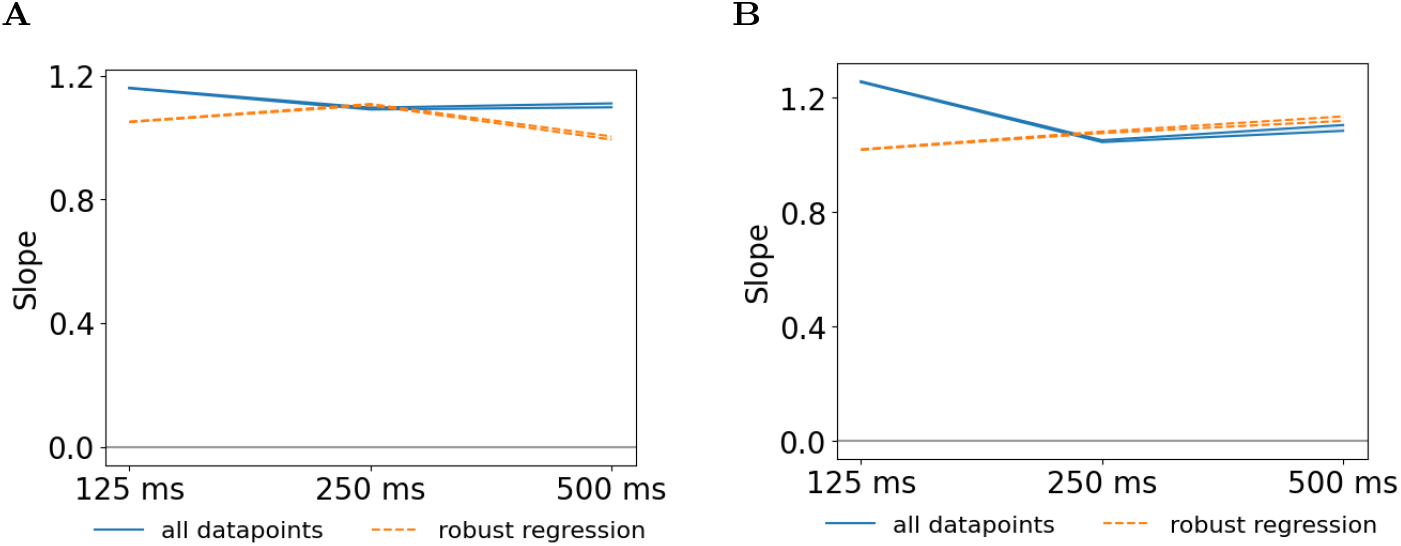
95% confidence intervals of slopes for NAD vs. TD of all points (blue solid) and robust regression (orange dashed) for 100 neurons by (A) 400 samples of spike rates and (B) 500 samples of spike rates with window lengths between 125ms to 500ms.

### 7.9 Details of likelihood ratio tests

The likelihood ratio tests compare the full model performance of TD+HS with reduced models TD-only or HS-only across NAD and sparseness measures. We randomly sampled equal numbers of data points across each cortical area and split them as 75%-25% to create a held-out validation set. The log-likelihood loss for the full model and reduced models was measured on the held-out set. We repeated this test 100 times and average the log-likelihood loss to compute the p-values. We report the variance-weighted mean of regression coefficients across these random draws.

#### 7.9.1 Neural activity dimension

**Table 1:** p-value for likelihood ratio test with TD representing values for only TD as the reduced model and HS values for only HS as the reduced model, and the full model is generated using both TD and HS to predict neural activity dimension. A p-value less than 0.05 indicates that the full model predicts neural activity dimension more effectively than the corresponding reduced model.

| window (ms) | TD + HS vs TD | TD + HS vs HS |
| --- | --- | --- |
| Linear Regression |  |  |
| 125 | 0.2028 | 0 |
| 250 | 1 | 0.0003 |
| 500 | 1 | 0.0066 |
| Robust Regression |  |  |
| 125 | 0.254 | 0 |
| 250 | 0.9166 | 0.0003 |
| 500 | 1 | 0.0066 |

**Table 2:** For neural activity dimension, full model columns TD and HS represent regression coefficient for TD and HS in the full model respectively for different window lengths. Reduced model column TD represents the regression coefficient for TD only model and column HS represents the regression coefficient for HS only model for different window lengths.

| window<br>(ms) | Full Model |  | Reduced Model |  |
| --- | --- | --- | --- | --- |
|  | (TD + HS) |  | (TD or HS) |  |
|  | TD | HS | TD | HS |
| Linear Regression |  |  |  |  |
| 125 | 1.265 | -2.234 | 1.098 | -0.224 |
| 250 | 1.184 | -1.711 | 1.057 | 0.229 |
| 500 | 1.274 | -2.159 | 1.114 | -0.100 |
| Robust Regression |  |  |  |  |
| 125 | 1.139 | -1.634 | 1.010 | 0.388 |
| 250 | 0.969 | -0.697 | 0.916 | 0.871 |
| 500 | 1.099 | -1.286 | 0.995 | 0.809 |

#### 7.9.2 Population sparseness

**Table 3:** p-value for likelihood ratio test with TD representing values for only TD as the reduced model and HS values for only HS as the reduced model, and the full model is generated using both TD and HS to predict population sparseness. A p-value less than 0.05 indicates that the full model predicts neural activity dimension more effectively than the corresponding reduced model.

| window (ms) | TD + HS vs TD |  | TD + HS vs HS |  |
| --- | --- | --- | --- | --- |
| | $S_K$ | $S_R$ | $S_K$ | $S_R$ |
| Linear Regression |  |  |  |  |
| 125 | $2.2 \times 10^{-187}$ | $2.4 \times 10^{-13}$ | 0.0 | 0.0 |
| 250 | $2.6 \times 10^{-77}$ | $1.5 \times 10^{-7}$ | 0.0 | 0.0 |
| 500 | $2.4 \times 10^{-62}$ | $7.9 \times 10^{-7}$ | $2.4 \times 10^{-265}$ | 0.0 |
| Robust Regression |  |  |  |  |
| 125 | $5.5 \times 10^{-111}$ | $1.4 \times 10^{-12}$ | 0.0 | 0.0 |
| 250 | $1.7 \times 10^{-45}$ | $6.2 \times 10^{-7}$ | 0.0 | 0.0 |
| 500 | $3.3 \times 10^{-37}$ | $5.3 \times 10^{-7}$ | $5.0 \times 10^{-220}$ | 0.0 |

**Table 4:** For population sparseness kurtosis (*S*_*K*_) and activity fraction (*S*_*R*_), full model columns TD and HS represent regression coefficients for TD and HS in the full model respectively for different window lengths. Reduced model column TD represents the regression coefficient for TD-only model and column HS represents the regression coefficient for HS-only model for different window lengths.

| window<br>(ms) | $S_K$ | | | | $S_R$ | | | |
| --- | --- | --- | --- | --- | --- | --- | --- | --- |
|  | Full Model<br>(TD + HS) |  | Reduced Model<br>(TD or HS) |  | Full Model<br>(TD + HS) |  | Reduced Model<br>(TD or HS) |  |
|  | TD | HS | TD | HS | TD | HS | TD | HS |
| Linear Regression |  |  |  |  |  |  |  |  |
| 125 | -2.54 | 9.29 | -1.85 | 5.17 | -0.04 | 0.02 | -0.03 | -0.04 |
| 250 | -2.40 | 8.64 | -1.75 | 4.77 | -0.04 | 0.02 | -0.03 | -0.04 |
| 500 | -2.59 | 11.57 | -1.73 | 7.38 | -0.03 | 0.02 | -0.03 | -0.03 |
| Robust Regression |  |  |  |  |  |  |  |  |
| 125 | -1.74 | 3.29 | -1.48 | 0.36 | -0.04 | 0.02 | -0.03 | -0.04 |
| 250 | -1.69 | 3.00 | -1.45 | 0.18 | -0.04 | 0.02 | -0.03 | -0.04 |
| 500 | -1.63 | 4.08 | -1.31 | 1.31 | -0.03 | 0.03 | -0.03 | -0.03 |

#### 7.9.3 Lifetime sparseness

**Table 5:** p-value for likelihood ratio test with TD representing values for only TD as the reduced model and HS values for only HS as the reduced model, and the full model generated using both TD and HS to predict lifetime sparseness. A p-value greater than 0.05 indicates that the reduced hypothesis may represent the lifetime sparseness as effectively as the full model.

| window (ms) | TD + HS vs TD |  | TD + HS vs HS |  |
| --- | --- | --- | --- | --- |
| | $S_K$ | $S_R$ | $S_K$ | $S_R$ |
| Linear Regression |  |  |  |  |
| 125 | 0.10 | $1.5 \times 10^{-3}$ | $2.5 \times 10^{-30}$ | $4.6 \times 10^{-119}$ |
| 250 | 0.09 | 0.14 | $2.7 \times 10^{-10}$ | $2.4 \times 10^{-52}$ |
| 500 | 0.06 | 0.82 | $2.2 \times 10^{-5}$ | $1.1 \times 10^{-28}$ |
| Robust Regression |  |  |  |  |
| 125 | 0.54 | $1.6 \times 10^{-3}$ | 1.0 | $6.0 \times 10^{-117}$ |
| 250 | 0.38 | 0.16 | 1.0 | $9.8 \times 10^{-52}$ |
| 500 | 0.22 | 1.00 | 0.14 | $1.7 \times 10^{-28}$ |

**Table 6:** For lifetime sparseness kurtosis (*S*_*K*_) and activity fraction (*S*_*R*_), full model columns TD and HS represent regression coefficients for TD and HS in the full model respectively for different window lengths. Reduced model column TD represents the regression coefficient for TD-only model and column HS represents the regression coefficient for HS-only model for different window lengths.

| window<br>(ms) | $S_K$ | | | | $S_R$ | | | |
| --- | --- | --- | --- | --- | --- | --- | --- | --- |
|  | Full Model<br>(TD + HS) |  | Reduced Model<br>(TD or HS) |  | Full Model<br>(TD + HS) |  | Reduced Model<br>(TD or HS) |  |
|  | TD | HS | TD | HS | TD | HS | TD | HS |
| Linear Regression |  |  |  |  |  |  |  |  |
| 125 | -0.54 | 0.74 | -0.49 | -0.13 | -0.03 | -0.04 | -0.03 | -0.09 |
| 250 | -0.63 | 1.80 | -0.49 | 0.76 | -0.03 | -0.03 | -0.03 | -0.07 |
| 500 | -0.87 | 3.85 | -0.59 | 2.41 | -0.03 | -0.01 | -0.03 | -0.06 |
| Robust Regression |  |  |  |  |  |  |  |  |
| 125 | -0.35 | 0.06 | -0.35 | -0.50 | -0.03 | -0.04 | -0.03 | -0.09 |
| 250 | -0.41 | 0.39 | -0.38 | -0.25 | -0.03 | -0.02 | -0.03 | -0.07 |
| 500 | -0.42 | 1.06 | -0.34 | 0.43 | -0.03 | -0.01 | -0.03 | -0.06 |

### 7.10 Approximating multidimensional cortical structure in convolutional networks

Convolutional networks often serve as abstract models of visual cortex. Despite various limitations, such as the biological implausibility of backpropagation (but see e.g. [70]), they have certain structural parallels and are strong predictors of brain activity [e.g., 55, 42]. The network structure of mouse visual cortex can be closely approximated by a suitably parameterized convolutional network [58]. We wondered whether convolutional networks could also approximate the higher-dimensional structure of multsensory convergence in the cortex. Qualitatively, we noticed that individual multisensory coordinates often do not vary linearly over space in higher areas, and that they have varied spatial frequencies (Section 7.5). To approximate these phenomena in a convolutional model, we used spatial remapping. Specifically, a multisensory layer receives differently-remapped convolutions from multiple unisensory layers,

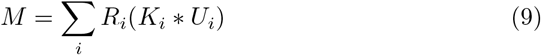

where *U*_*i*_ are unisensory layers, *K*_*i*_ are corresponding convolution kernels, and *R*_*i*_ are corresponding remapping functions. The remapping functions move pixels to new coordinates, interpolating as necessary to allow local stretching, compression, etc. of the original representations.

Figure 21 illustrates this model. Panel A shows an example of a remapping function with a certain spatial frequency distribution. Panel B shows how the dimension of the multisensory map varies with the spatial frequency content and the correlations between remapping functions. The resulting dimension is higher when the remapping functions have higher spatial frequencies, and lower when the remapping functions are correlated. With two-dimensional input from six different modalities, the dimension can range from two to twelve depending on these parameters. Differences in connection weights would also reduce TD in this model. Alternative models may also account for differences in spatial frequency and dimension.

**Figure 21:**
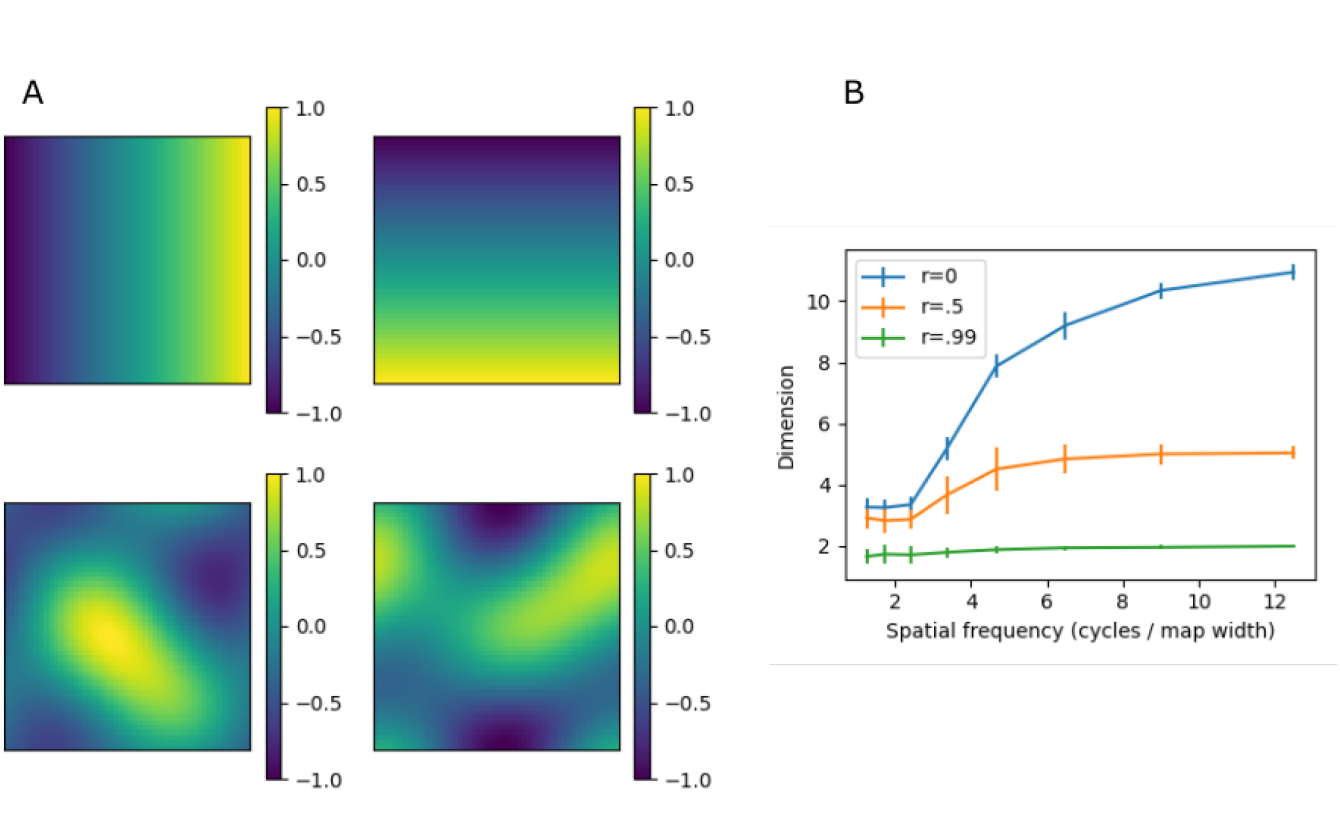
(A) Example remapping with frequencies up to 2.5 cycles across the feature map. The top-left and top-right images show raw (unmapped) horizontal and vertical coordinates. The images below show the same coordinates remapped to different positions. (B) Topography dimension (TD) of convolutional layers that employ such remappings. The layers receive inputs with two-dimensional topographies from six modalities, and each input is spatially remapped. TD depends on the bandwidths of the remapping functions, and correlations between different modalities’ remapping functions. Error bars show mean +/-standard deviation over 25 simulations with random remapping functions. For these simulations, remapping functions were first generated as whitenoise images. Spatial frequency content was controlled via low-pass filtering with cutoff frequencies as indicated. To introduce correlations, the remapping functions of different modalities were organized into pixel-wise vectors. Then these vectors were multiplied by a matrix *L*, where *LL*^*T*^ = *A, A* is a correlation matrix with *r* in the off-diagonal entries, and *L* is from the Cholesky decomposition.

## References

[1] Josh Achiam et al. “Gpt-4 technical report”. In: arXiv preprint 2303.08774 (2023).

[2] Annette E Allen et al. “Convergence of visual and whisker responses in the primary somatosensory thalamus (ventral posterior medial region) of the mouse”. In: The Journal of physiology 595.3 (2017), pp.865–881.

[3] Michael Beyeler et al. “Neural correlates of sparse coding and dimension-ality reduction”. In: PLoS computational biology 15.6 (2019), e1006908.

[4] Malte Bieler et al. “Multisensory integration in rodent tactile but not visual thalamus”. In: Scientific Reports 8.1 (2018), p.15684.

[5] Yinan Cao et al. “Causal inference in the multisensory brain”. In: Neuron 102.5 (2019), pp.1076–1087.

[6] Charlotte Caucheteux, Alexandre Gramfort, and Jean-Rémi King. “Evidence of a predictive coding hierarchy in the human brain listening to speech”. In: Nature human behaviour 7.3 (2023), pp.430–441.

[7] Theresa Cooke et al. “Multimodal similarity and categorization of novel, three-dimensional objects”. In: Neuropsychologia 45 (2007), pp.484–495. doi: 10.1016/j.neuropsychologia.2006.02.009.

[8] Ivan E De Araujo and Sidney A Simon. “The gustatory cortex and multisensory integration”. In: International journal of obesity 33.2 (2009), S34– S43.

[9] Frédérique De Vignemont. “Multimodal unity and multimodal binding”. In: David Bennett and Christopher Hill. Sensory integration and the unity of consciousness. MIT Press, 2014, pp.125–150.

[10] Jean-René Duhamel Carol L Colby, and Michael E Goldberg. “Ventral intraparietal area of the macaque: congruent visual and somatic response properties”. In: Journal of neurophysiology 79.1 (1998), pp.126–136.

[11] Marc O Ernst and Martin S Banks. “Humans integrate visual and haptic information in a statistically optimal fashion”. In: Nature 415.January (2002), pp.429–433.

[12] Daniel J Felleman and David C Van Essen. “Distributed hierarchical processing in the primate cerebral cortex.” In: Cerebral cortex (New York, NY: 1991) 1.1 (1991), pp.1–47.

[13] Christopher R Fetsch et al. “Neural correlates of reliability-based cue weighting during multisensory integration”. In: Nature Neuroscience 15.1 (2012). doi: 10.1038/nn.2983.

[14] Peiran Gao et al. “A theory of multineuronal dimensionality, dynamics and measurement”. In: BioRxiv (2017), p.214262.

[15] Aleena R Garner and Georg B Keller. “A cortical circuit for audio-visual predictions”. In: Nature neuroscience 25.1 (2022), pp.98–105.

[16] Richard C Gerum et al. “Different spectral representations in optimized artificial neural networks and brains”. In: arXiv preprint 2208.10576 (2022).

[17] Asif A Ghazanfar and Charles E Schroeder. “Is neocortex essentially multisensory?” In: Trends in cognitive sciences 10.6 (2006), pp.278–285.

[18] Sara R J Gilissen et al. “Reconsidering the border between the visual and posterior parietal cortex of mice”. In: Cerebral Cortex 31 (2021), pp.1675– 1692. doi: 10.1093/cercor/bhaa318.

[19] Lindsey L Glickfeld and Shawn R Olsen. “Higher-order areas of the mouse visual cortex”. In: Annual review of vision science 3.1 (2017), pp.251–273.

[20] Andrea M Green and Dora E Angelaki. “Multisensory integration: resolving sensory ambiguities to build novel representations”. In: Current opinion in neurobiology 20.3 (2010), pp.353–360.

[21] Julie A Harris et al. “Hierarchical organization of cortical and thalamic connectivity”. In: Nature 575.7781 (2019), pp.195–202.

[22] Ruey-Song Huang et al. “Mapping multisensory parietal face and body areas in humans”. In: Proceedings of the National Academy of Sciences 109.44 (2012), pp.18114–18119.

[23] Peter J. Huber. “Robust Estimation of a Location Parameter”. In: The Annals of Mathematical Statistics 35.1 (1964), pp.73–101. doi: 10.1214/aoms/1177703732. url: http://doi.org/10.1214/aoms/1177703732.

[24] Howard C Hughes et al. “Visual-auditory interactions in sensorimotor processing: Saccades versus manual responses”. In: Journal of Experimental Psychology. Human Perception and Performance 20.1 (1994), pp.131– 153.

[25] Saad Jbabdi, Stamatios N Sotiropoulos, and Timothy E Behrens. “The topographic connectome”. In: Current Opinion in Neurobiology 23 (2012), pp.207–215. issn: 0959-4388. doi: 10.1016/j.conb.2012.12.004. url: http://dx.doi.org/10.1016/j.conb.2012.12.004.

[26] Stephen E Jones, Bradley R Buchbinder, and Itzhak Aharon. “Three-dimensional mapping of cortical thickness using Laplace’s equation”. In: Human brain mapping 11.1 (2000), pp.12–32.

[27] Salman Khan, Alexander Wong, and Bryan Tripp. “Modeling the Role of Contour Integration in Visual Inference”. In: Neural Computation 36.1 (2023), pp.33–74.

[28] Meenakshi Khosla et al. “Cortical response to naturalistic stimuli is largely predictable with deep neural networks”. In: Science Advances 7.22 (2021), eabe7547.

[29] Thomas Knöpfel et al. “Audio-visual experience strengthens multisensory assemblies in adult mouse visual cortex”. In: Nature communications 10.1 (2019), p.5684.

[30] Joseph E Knox et al. “High-resolution data-driven model of the mouse connectome”. In: Network Neuroscience 3.1 (2018), pp.217–236.

[31] Konrad P Körding et al. “Causal inference in multisensory perception”. In: PLoS one 2.9 (2007), e943.

[32] International Brain Laboratory et al. “A brain-wide map of neural activity during complex behaviour”. In: Biorxiv (2023), pp.2023–07.

[33] Dmitry Lyamzin and Andrea Benucci. “The mouse posterior parietal cortex: Anatomy and functions”. In: Neuroscience research 140 (2019), pp.14– 22.

[34] Harry McGurk and John MacDonald. “Hearing lips and seeing voices”. In: Nature 264.5588 (1976), pp.746–748.

[35] Guido T Meijer et al. “Audiovisual integration enhances stimulus detection performance in mice”. In: Frontiers in Behavioral Neuroscience 12.October (2018), pp.1–12. doi: 10.3389/fnbeh.2018.00231.

[36] Jonathan A Michaels et al. “A goal-driven modular neural network predicts parietofrontal neural dynamics during grasping”. In: Proceedings of the national academy of sciences 117.50 (2020), pp.32124–32135.

[37] Anna S Mitchell et al. “Retrosplenial cortex and its role in spatial cognition”. In: Brain and neuroscience advances 2 (2018), p.2398212818757098.

[38] Mai M Morimoto, Emi Uchishiba, and Aman B Saleem. “Organization of feedback projections to mouse primary visual cortex”. In: IScience 24.5 (2021).

[39] X Ryan J Morrill and X Andrea R Hasenstaub. “Visual information present in infragranular layers of mouse auditory cortex”. In: Journal of Neuroscience 38.11 (2018), pp.2854–2862. doi: 10.1523/JNEUROSCI.3102-17.2018.

[40] Mouse Connectivity Models. https://github.com/AllenInstitute/mouse_connectivity_models. 2018.

[41] Micah M Murray et al. “The multisensory function of the human primary visual cortex”. In: Neuropsychologia 83 (2016), pp.161–169.

[42] Aran Nayebi et al. “Mouse visual cortex as a limited resource system that self-learns an ecologically-general representation”. In: PLOS Computational Biology 19.10 (2023), e1011506.

[43] Uta Noppeney. “Perceptual inference, learning, and attention in a multisensory world”. In: Annual review of neuroscience 44 (2021), pp.449– 473.

[44] Seung Wook Oh et al. “A mesoscale connectome of the mouse brain”. In: Nature 508.7495 (2014), pp.207–214.

[45] Umberto Olcese, Giuliano Iurilli, and Paolo Medini. “Cellular and synaptic architecture of multisensory integration in the mouse neocortex”. In: Neuron 79.3 (2013), pp.579–593.

[46] Matthijs N Oude Lohuis et al. “Triple dissociation of visual, auditory and motor processing in mouse primary visual cortex”. In: Nature Neuroscience 27.4 (2024), pp.758–771.

[47] Andrew Owens and Alexei A. Efros. “Audio-Visual Scene Analysis with Self-Supervised Multisensory Features”. In: Proceedings of the European Conference on Computer Vision (ECCV). Sept. 2018.

[48] Karl Pearson. ““The error law and its generalizations by Fechner and Pearson.” a rejoinder”. In: Biometrika 4.1-2 (1905), pp.169–212.

[49] Alexandre Pouget, Sophie Deneve, and Jean-René Duhamel. “A computational perspective on the neural basis of multisensory spatial representations”. In: Nature Reviews Neuroscience 3.9 (2002), pp.741–747.

[50] Alec Radford et al. “Learning Transferable Visual Models From Natural Language Supervision”. In: Proceedings of the 38th International Conference on Machine Learning. Ed. by Marina Meila and Tong Zhang. Vol. 139. Proceedings of Machine Learning Research. PMLR, 18–24 Jul 2021, pp.8748–8763. url: https://proceedings.mlr.press/v139/radford21a.html.

[51] Stefano Recanatesi et al. “A scale-dependent measure of system dimensionality”. In: Patterns 3.8 (2022).

[52] Ramon Reig and Gilad Silberberg. “Multisensory integration in the mouse striatum”. In: Neuron 83.5 (2014), pp.1200–1212.

[53] Omid Rezai, Lucas Stoffl, and Bryan Tripp. “How are response properties in the middle temporal area related to inference on visual motion patterns?” In: Neural Networks 121 (2020), pp.122–131.

[54] Edmund T Rolls and Martin J Tovee. “Sparseness of the neuronal representation of stimuli in the primate temporal visual cortex”. In: Journal of neurophysiology 73.2 (1995), pp.713–726.

[55] Martin Schrimpf et al. “Brain-score: Which artificial neural network for object recognition is most brain-like?” In: BioRxiv (2018), p.407007.

[56] Martin I Sereno and Ruey-Song Huang. “Multisensory maps in parietal cortex”. In: Current Opinion in Neurobiology 24 (2014), pp.39–46. issn: 0959-4388. doi: 10.1016/j.conb.2013.08.014. url: http://dx.doi.org/10.1016/j.conb.2013.08.014.

[57] Kamal Shadi, Eva Dyer, and Constantine Dovrolis. “Multisensory integration in the mouse cortical connectome using a network diffusion model”. In: Network Neuroscience 4.4 (2020), pp.1030–1054.

[58] Jianghong Shi et al. “MouseNet: A biologically constrained convolutional neural network model for the mouse visual cortex”. In: PLOS Computational Biology 18.9 (2022), e1010427.

[59] Joshua H Siegle et al. “Survey of spiking in the mouse visual system reveals functional hierarchy”. In: Nature 592.7852 (2021), pp.86–92.

[60] Martin Stacho and Denise Manahan-Vaughan. “Mechanistic flexibility of the retrosplenial cortex enables its contribution to spatial cognition”. In: Trends in neurosciences 45.4 (2022), pp.284–296.

[61] Barry E Stein, Terrence R Stanford, and Benjamin A Rowland. “Development of multisensory integration from the perspective of the individual neuron”. In: Nature Reviews Neuroscience 15.August (2014). issn: 1471-003X. doi: 10.1038/nrn3742.

[62] Nicholas A Steinmetz et al. “Distributed coding of choice, action and engagement across the mouse brain”. In: Nature 576.7786 (2019), pp.266– 273.

[63] Larry W Swanson, Joel D Hahn, and Olaf Sporns. “Organizing principles for the cerebral cortex network of commissural and association connections”. In: Proceedings of the National Academy of Sciences 114.45 (2017), E9692–E9701.

[64] Tsung-Chih Tsai et al. “Distinct contribution of granular and agranular subdivisions of the retrosplenial cortex to remote contextual fear memory retrieval”. In: Journal of Neuroscience 42.5 (2022), pp.877–893.

[65] Alice B Van Derveer, Jordan M Ross, and Jordan P Hamm. “Robust multisensory deviance detection in the mouse parietal associative area”. In: Current Biology 33.18 (2023), pp.3969–3976.

[66] William E Vinje and Jack L Gallant. “Sparse coding and decorrelation in primary visual cortex during natural vision”. In: Science 287.5456 (2000), pp.1273–1276.

[67] Mark T Wallace, M Alex Meredith, and Barry E Stein. “Multisensory integration in the superior colliculus of the alert cat”. In: Journal of neurophysiology 80.2 (1998), pp.1006–1010.

[68] Quanxin Wang et al. “The Allen mouse brain common coordinate framework: a 3D reference atlas”. In: Cell 181.4 (2020), pp.936–953.

[69] Johan Winnubst et al. “Reconstruction of 1,000 projection neurons reveals new cell types and organization of long-range connectivity in the mouse brain”. In: Cell 179.1 (2019), pp.268–281.

[70] Will Xiao et al. “Biologically-plausible learning algorithms can scale to large datasets”. In: arXiv preprint 1811.03567 (2018).

[71] Xiaohua Zhai et al. “Scaling vision transformers”. In: Proceedings of the IEEE/CVF conference on computer vision and pattern recognition. 2022, pp.12104–12113.

